# Decoding movie content from neuronal population activity in the human medial temporal lobe

**DOI:** 10.1101/2024.06.13.598791

**Authors:** Franziska Gerken, Alana Darcher, Pedro J Gonçalves, Rachel Rapp, Ismail Elezi, Johannes Niediek, Marcel S Kehl, Thomas P Reber, Stefanie Liebe, Jakob H Macke, Florian Mormann, Laura Leal-Taixé

**Affiliations:** Dynamic Vision and Learning Group, Technical University of Munich, Munich, Germany; Dept. of Epileptology, University Medical Center of Bonn, Bonn, Germany; Machine Learning in Science, Excellence Cluster Machine Learning and Tübingen AI Center, University of Tübingen, Tübingen, Germany; VIB-Neuroelectronics Research Flanders (NERF), Belgium; imec, Belgium; Department of Epileptology and Neurology, University Hospital Tübingen, Tübingen, Germany; Empirical Inference, Max Planck Institute for Intelligent Systems, Tübingen, Germany; Machine Learning Group, Technical University of Berlin, Berlin, Germany; Department of Experimental Psychology, University of Oxford, Oxford, UK; Faculty of Psychology, UniDistance Suisse, Brig, Switzerland

## Abstract

The human medial temporal lobe (MTL), a region implicated in memory and high-level cognition, contains neurons that respond selectively to stimuli belonging to specific categories, such as individual people, landmarks, or objects. However, these neurons have been largely studied via static, isolated presentations of stimuli. Therefore, it is unclear how neurons in the MTL respond to rich stimuli such as movies, and which dynamical stimulus features can be retrieved from neuronal population spiking activity. We studied single-unit responses from 2286 neurons recorded from the amygdala, hippocampus, entorhinal cortex, and parahippocampal cortex of 29 intracranially implanted patients during the presentation of an 83-minute movie. We found only a few individual neurons that exhibited a classic selective response to semantic features. However, we successfully decoded the presence of characters, settings, and visual transitions from neuronal population activity. The information relevant for decoding varies across regions depending on the feature category, as visual transitions could be decoded from subsets of neurons with selective responses, whereas character and location features relied on distributed representations. Our results demonstrate an approach for reliably decoding movie features in the human MTL, and suggest that the brain uses a population code when representing character and location features.

## Introduction

The human medial temporal lobe (MTL) plays an integral role in the representation of semantic information. Single neurons in the MTL exhibit strong and highly selective tunings to categories, such as faces or locations ^1,2^, and to specific concepts, such as individual celebrities or objects ^3,4^. These semantically tuned neurons relate to the processing of information at a conscious, declarative level, as their activity varies depending on perception ^5–7^—when images are presented but unseen, these neurons exhibit reduced and delayed spiking compared to consciously seen images ^8^. Such cells also support the formation of new memories ^9^, and are involved in the retrieval of previously encoded experiences ^10,11^.

Studies investigating the representation of semantic information in the human MTL have mostly focused on characterizing these neurons individually, often without considering their population dynamics. Many of these studies have identified semantically tuned cells by screening for stimulus-selective responses to static images depicting isolated persons or objects ^12,13^. While this approach is effective for probing specific functional properties of individual neurons, it limits the generalizability of findings to more complex and dynamic contexts. Naturalistic and dynamic stimuli, such as movies, provide closer approximations to real-world environments, but also pose substantial challenges, as they are presented continuously and depict complex evolving scenes. Several studies have used functional magnetic resonance imaging (fMRI) to study neural responses to natural movies. One line of work has focused on representations in early visual areas ^14,15^ and across the cortex ^16,17^. Another line of work has identified synchrony in the brain states of separate individuals viewing a common movie ^18,19^. Such states appear to be particularly well aligned in individual brains surrounding transition events within an ongoing continuous stimulus, as reported in fMRI studies ^20,21^, intracranial field potentials along the human ventral visual pathway ^22^, and in single neurons in the human parahippocampal gyrus, hippocampus, and amygdala ^11^. Most single neuron studies which use movies as stimuli have been based on only short ‘snippets’ of movies, and have so far largely examined representation on the level of the individual neuron ^23^ with few having addressed frame-wise representations in longer movie sequences ^24^. However, it remains an open question of how populations of neurons in the MTL collectively respond to naturalistic visual stimuli, such as those encountered in real-world environments, and which features of population activity encode information about specific components of such stimuli.

In this study, we investigate how information embedded in a naturalistic and dynamic stimulus is processed by neuronal populations in the human medial temporal lobe (MTL). Specifically, we asked the following questions: (1) Which aspects of a movie’s content can be decoded from neuronal activity in the MTL? (2) Which brain regions are informative for specific stimulus categories (e.g. visual transitions or characters)? (3) Is the relevant information distributed across the neuronal population? We recorded the activity of neurons from patients with intracranially implanted electrodes as each watched the full-length commercial movie “500 Days of Summer”. Our dataset is unique in both size and duration: we recorded a total of 2286 neurons across 29 patients during the complete presentation of the movie. To analyze the relationship between the neuronal activity and the film’s content, we labeled the presence of main characters, whether a scene was indoors or outdoors, and visual transitions of the movie on a frame-by-frame basis.

We introduce a machine learning-based decoding pipeline that decodes a movie’s visual content from population-level neuronal activity. At the single-unit level, individual neurons generally lacked reliable responses to the visual features, and we observed consistent stimulus-related changes in firing rates primarily during visual transitions. However, at the population level, we achieved strong decoding performance across all visual features. For visual transitions, neurons exhibiting consistent changes in activity played a key role in population-level decoding. In contrast, no similar pattern emerged when decoding character identities. By analyzing the contributions of individual neurons, we identified distinct subsets of neurons that influenced decoding performance, extending beyond the subsets identified by stimulus-aligned changes in firing activity. Remarkably, we found that restricting the analysis to a substantially smaller subset of these key neurons was sufficient to replicate the full population-level decoding performance. Taken together, our findings show that information about dynamic stimuli can be decoded from neuronal population activity, even in the absence of strong single-neuron selectivity.

## Results

We recorded from 29 patients (17 female, ages 22 *→* 63) as each watched the movie *500 Days of Summer* (83 minutes) in its entirety (Fig. 1a). Patients were bilaterally implanted with depth electrodes for seizure monitoring, and spiking activity was recorded from a total of 2286 single- and multi-neurons across the amygdala (A; 580 neurons, 25.37%), hippocampus (H; 794, 34.73%), entorhinal cortex (EC; 440, 19.25%), parahippocampal cortex (PHC; 373, 16.32%), and piriform cortex (PIC; 99, 4.33%) (Fig. 1b). We pooled the neurons across patients (distribution shown in Supp. Fig. S1) and performed subsequent analyses on the resulting population. Due to the low number of neurons relative to the complete population, the PIC was excluded from subsequent region-wise analyses.

**Figure 1.**
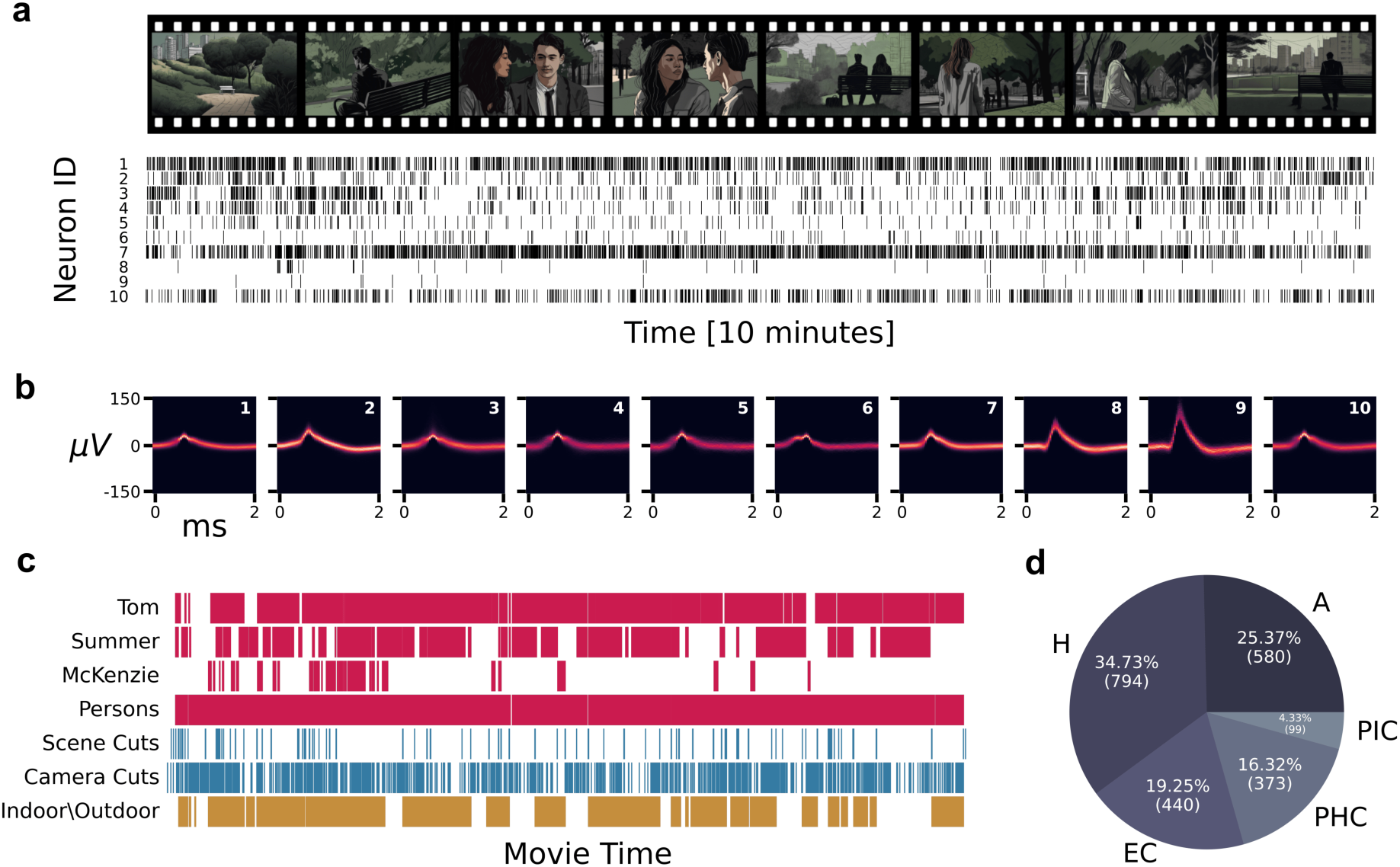
Overview of dataset, features, and decoding approach a) Example of the recorded neuronal activity. Patients watched the complete commercial film *500 Days of Summer* while neuronal activity was recorded via depth electrodes. Top row: example movie frames. Due to copyright, the original movie frames have been replaced with images generated using stable diffusion ^25^. Bottom row: spike trains from ten amygdala neurons of a single patient, where each row shows data from an individual neuron (corresponding ID number given as a label). b) Spike density plot showing the waveforms of each neuron in a (corresponding neuron ID given in top right). Neurons shown include both single- and multi-neurons. c) Distribution of labels across the entire movie (runtime: 83 minutes). Occurrences of character-related features are in magenta, visual transition events in blue, and location events in yellow. d) Distribution of the 2286 neurons across the recorded regions (A: amygdala, H: hippocampus, EC: entorhinal cortex, PHC: parahippocampal cortex, PIC: piriform cortex) for all 29 patients.

To determine which features of the movie are represented in MTL activity, we obtained frame-wise annotations of character presence, indoor/outdoor setting, and the occurrence of a visual transition, i.e. camera cuts and scene cuts (similar to “soft” and “hard” boundaries ^11^) (Fig. 1c). All annotations concern the visual occurrence of a given feature in the frame, and do not consider feature content of the movie’s audio. We used a mixture of manual and automated methods for obtaining these annotations (see Methods; sample frames in Supp. Fig. S2). For character-related content, we restricted the analysis to those characters most relevant to the movie’s narrative (Tom, Summer, McKenzie) as determined by screentime. For the analysis of location, we investigate indoor and outdoor settings. Note that these two features are mutually exclusive, and the annotation of both are combined into a single label.

### Stimulus-aligned responsive neurons found primarily in parahippocampal cortex

To investigate whether the visual features of the movie are encoded at the level of individual neurons, we analyzed stimulus-evoked changes in the firing rate of neurons after the onset of characters, indoor and outdoor scenes, and visual transitions, for each MTL subregion separately (Fig. 2a, Supp. Fig. S3 - S5). We classified each individual neuron as responsive or non-responsive according to a previously established criterion ^8^ which compares the spiking activity after stimulus onset to that of a baseline period, adapted to the dynamic presentation. Specifically, for each annotated feature, we identified all instances where the feature appeared following at least 1000 ms of absence and remained continuously present for at least 1000 ms. We applied a cluster permutation test ^26^ to find time-points in which the firing rate between the sets of responsive and non-responsive neurons differ (see Methods for additional details). We found individual neurons with significant stimulus-evoked responses in all regions for visual transitions, Persons, and the characters Tom and Summer, including Face-only onsets (bin-wise Wilcoxon signed-rank test, Simes-corrected, *ω* = 0.001, see Supp. Table S1). Of these stimulus-evoked changes, neurons in the parahippocampal cortex responded to the largest set of stimulus features (Summer, Persons, Camera Cuts, Scene Cuts, Outdoor; *p ↑* 0.001, cluster permutation test), as compared to the hippocampus (Tom, Camera Cuts, Outdoor), amygdala (Camera Cuts), and entorhinal cortex (None). Over half of the parahippocampal neurons responded to Camera Cuts during the movie (193*/*373, 51.74%, bin-wise Wilcoxon signed-rank test, Simes-corrected, *ω* = 0.001) (Fig. 2b), and stimulus-evoked changes in firing were consistent across the set of responsive neurons (*p ↑* 0.001, cluster permutation test). For the remaining regions, stimulus-evoked modulations were less consistent across individual neurons (see Supp. Fig. S5a). Comparatively few parahippocampal neurons responded to Scene Cuts (25*/*373, 6.70%), although there was nonetheless a consistent pattern of modulation across responsive neurons (*p* = 0.018, cluster permutation test) (Fig. 2b). A similar pattern was observed for the onset of Outdoor scenes in the parahippocampal cortex (4*/*373, 1.07%). No responses were detected for the onset of indoor scenes. For characters, significant stimulus-evoked modulation in the responsive neurons were only observed for Summer (PHC, *p* = 0.006, cluster permutation test) and, to a lesser extent, Tom (H, *p* = 0.022, cluster permutation test) (Fig. 2a, Supp. Fig. S3). Taken together, the parahippocampal cortex contained neuronal subsets that showed a consistent pattern of increased firing after the onset of a visual transition (Camera or Scene Cut) within the movie. Characters and character faces evoked a clear change in firing activity at the subpopulation-level in the parahippocampal cortex, and sparingly in the hippocampus, but not in other tested regions.

**Figure 2.**
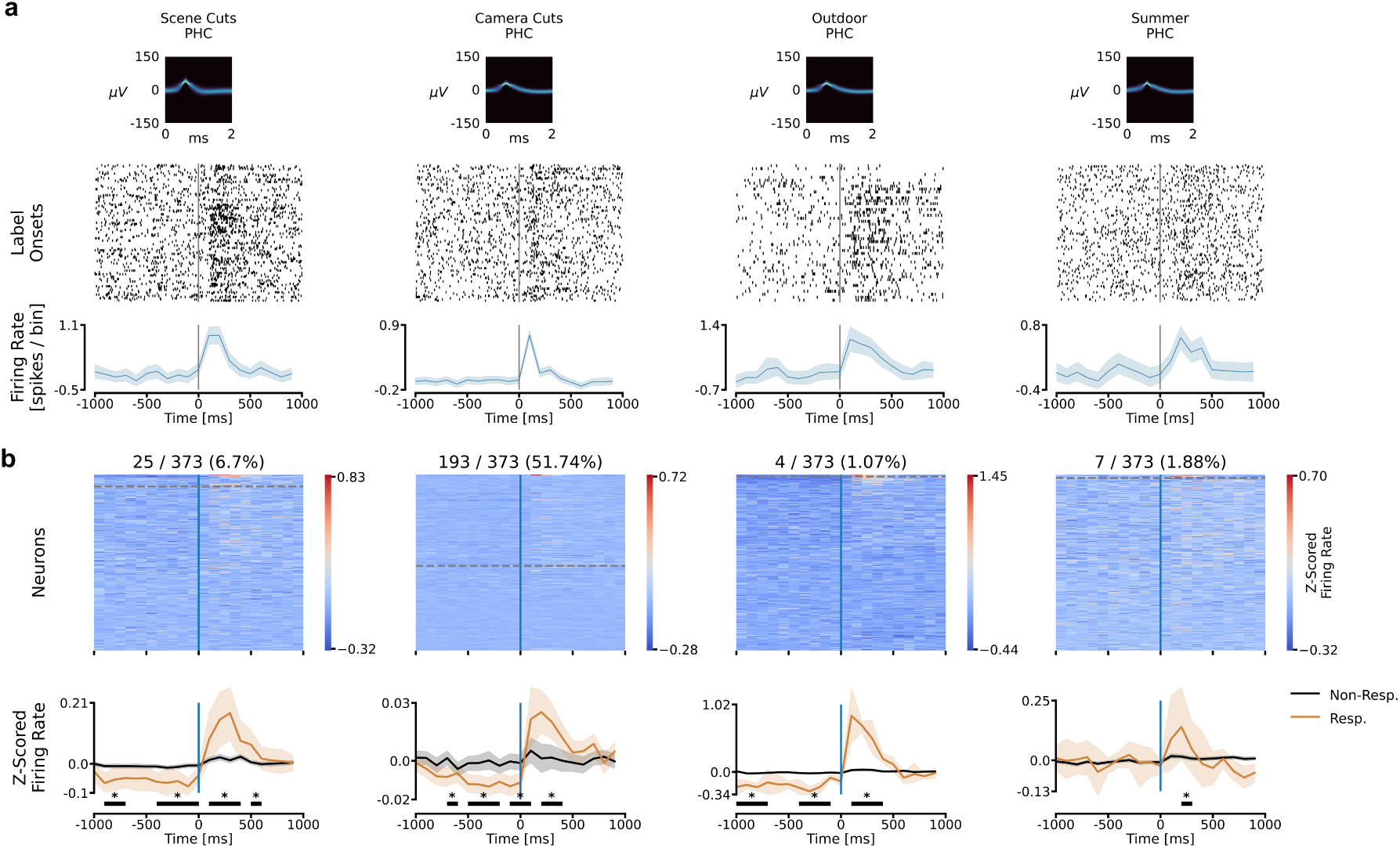
Responsive single-neurons in the parahippocampal cortex a) Example peri-stimulus activity for representative parahippocampal (PHC) neurons, for labels with a significant PHC response. Upper plots: spike density plot showing the waveforms a given responsive neuron (label name given as title). Middle plots: spike time rasters showing the neuron’s activity surrounding the onset of the corresponding label throughout the movie. Note: onsets for Scene Cuts and Camera Cuts were randomly subsampled to match the number of Summer appearances. Lower plots: average firing rate across 100 ms bins, for 1000 ms before and 1000 ms after the onset or event. Solid lines show the mean across all neurons, within group, and the transparent area shows the 95% confidence interval. b) Region-wise single-neuron activity surrounding the onset of labeled entity. Number of cells exhibiting a significant response over the total number of PHC cells are given as the title, followed by the corresponding percentage. Upper plots (heatmaps): averages of peri-stimulus spike rates per neuron (spikes per 100 ms bin, z-scored across the pseudotrial) for 1000 ms before and 1000 ms after label onset. Each row of the heatmap represents the average binned activity for one neuron. Neurons are sorted in descending order by the p-value of the response—the dotted grey line shows the threshold for responsive neurons (p *→* 0.001). Lower plots (line plots): average z-scored firing rate across bins. Neurons are separated into responsive (orange line) and non-responsive (black line). Solid lines show the mean for each group of neurons (responsive vs. non-responsive) and the transparent area depicts the 95% confidence interval. Significant differences between the responsive and non-responsive firing rates are shown as solid black lines (*, p *→* 0.05, cluster permutation test).

### Decoding of semantic content from population responses

The observed pattern of responses in individual neurons aligned to feature onsets suggests that these cells primarily carry information relating to visual transitions, and, to a lesser degree, the characters Summer and Tom, and Persons. However, previous work suggests that there may be differences in the coding capacity at the single-neuron and population level ^27^. Could there be stimulus-related information represented at the population level that is not apparent in the responses of individual neurons? To explore this, we decoded character presence, location, and visual transitions from the aggregated activity of the entire neuronal population using a recurrent neural network designed to capture both within-neuron and between-neuron activity patterns.

#### Decoding from neuronal activity

We aligned the activity of neurons across patients using the movie frames as a common time reference, generating a single neuronal pseudo-population, as done in previous work decoding from populations of single neurons ^28,29^. We first evaluated single-neuron decoding performance for the main character, Summer, by fine-tuning a firing activity threshold for each neuron. Using the total spike count around frame onset (spanning 800 ms before and after onset), we optimized a threshold to the validation set from each cell and subsequently predicted Summer’s appearance in the test set. This established a performance baseline for decoding character presence solely from the firing of individual neurons, without machine learning-based decoding algorithms. Decoding performances obtained by simply applying a threshold to single-neuron activity did not reliably predict the content, as the majority of neurons in the population performed near chance level (Fig. 3b). We then extended this analysis to a population-based approach, employing a Long Short-Term Memory (LSTM) network ^30^, a deep neural network well-suited for processing dynamic sequential data (Fig. 3a). This population-based approach improved decoding performance for Summer, surpassing the performances achieved by individual neurons alone. Decoding performances for all tested labels significantly exceeded chance level (zero for Cohen’s Kappa), as shown in Fig. 3c. The Persons label achieved the highest performance (mean *±* standard error of the mean, SEM: 0.36 *±* 0.05), followed by the location-related label Indoor/Outdoor (0.31 *±* 0.06). Note that in contrast to the single unit analysis, there is no need to conduct two separate trainings for the Indoor/Outdoor label since the label combines both features and the network implicitly learns to differentiate between the two. Character-specific labels (Summer, Tom, McKenzie) showed comparable performances, ranging between 0.23 and 0.32. Labels related to visual transitions in the movie exhibited the lowest but nonetheless significant performances of 0.2 *±* 0.03 and 0.18 *±* 0.01, respectively. Decoding results were consistent across metrics (F1 Score, PR-AUC, and AUROC shown in Supp. Table S2). We evaluated the statistical significance of our results by randomly permuting the test set labels (*N* = 1000), demonstrating that all decoding performances were significantly above chance level at an alpha level of 0.001 (see Methods). To ensure that this significance was not an artifact of the permutation procedure, we conducted an additional test by circularly shifting the neuronal data relative to the movie features (see Methods for more detail). This approach preserved the temporal structure of each dataset while disrupting the relationship between the neuronal data and the stimulus. Results were consistent between random permutation and circular shifts (Supp. Fig. S11).

**Figure 3.**
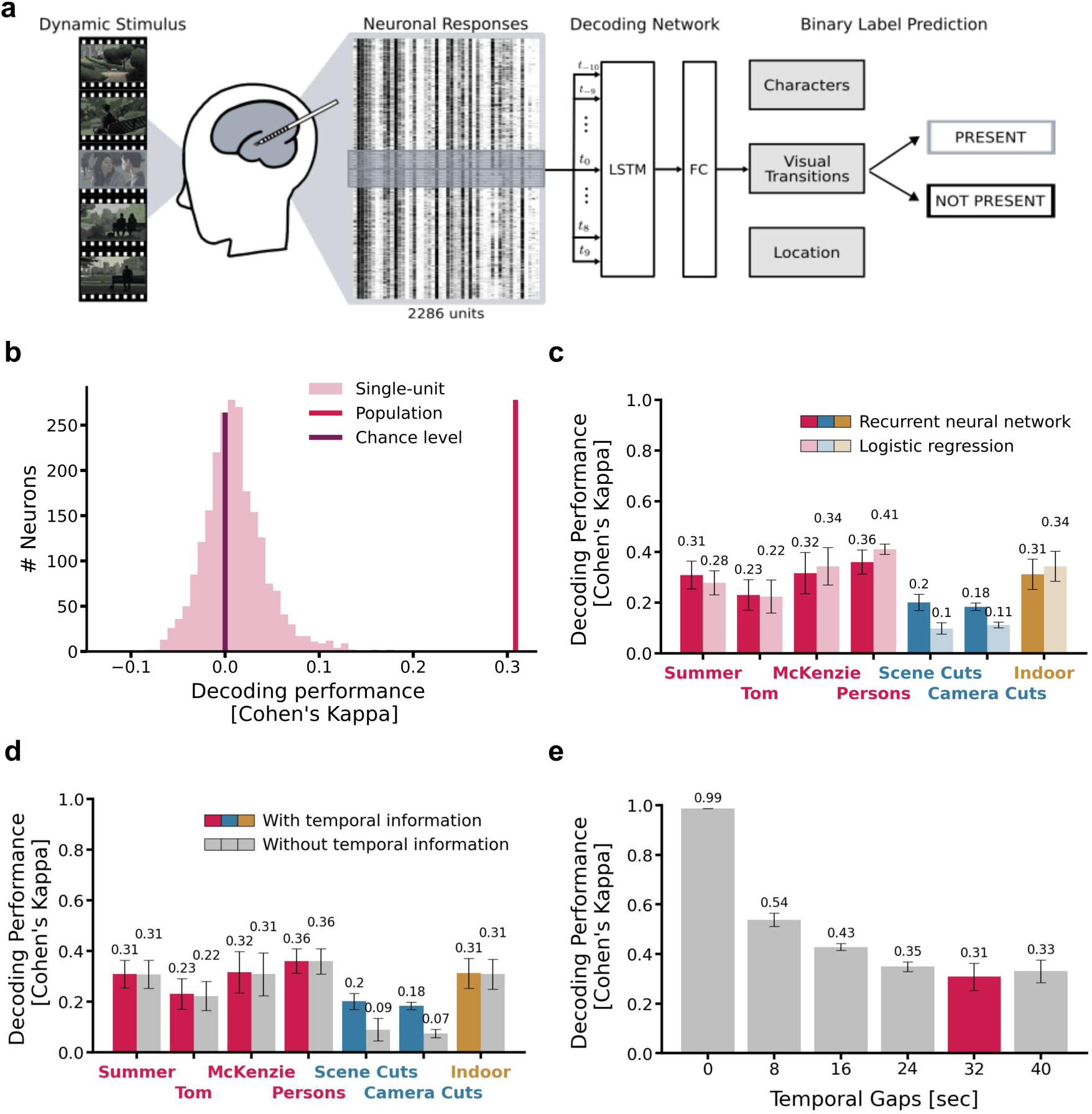
Categories of labels can be decoded from the neuronal population activity a) Overview of the neuronal decoding pipeline. Spiking data (individual neurons shown as columns) was sectioned into 1600 ms sequences, 800 ms before and after a frame onset (purple highlight; bins shown as horizontal lines, not to scale), and given as input to a two-layer Long Short-Term Memory (LSTM) network. The output of the fully connected layer (FC) predicts the presence of a given label in a frame. b) Assessment of individual-neuron decoding performance by classifying data samples into positive or negative predictions for the label Summer based on the firing activity of a neuron. c) Decoding performances on labels of the movie (reported performance using Cohen’s Kappa, mean performance across five different data splits, error bars indicate standard error of the mean). Labels fall into one of three categories—characters (pink), visual transitions (blue), or location (yellow)—with a separate model for each label. All performances were significantly better than chance level with an alpha level of 0.001. Decoding performances for the logistic regression model in lighter colors. d) Impact of temporal information in spike trains for recurrent neural networks. Trained models were evaluated using temporally altered test data (sequence order shuned, repeated 100 times). Colored bars depict performance without shuning, while grey bars represent shuned scenarios (reported performance using Cohen’s Kappa, results show the mean performance across five different data splits, variance given as standard error of the mean). e) Decoding performance of the main movie character (Summer) for different temporal gap sizes between samples of training, validation, and test sets. Colored temporal gap of 32 s indicates the chosen gap size for all reported performances in this study.

To further test the hypothesis that decoding is based on population activity rather than individual neurons, we also trained an LSTM network on individual neurons. From the 46 neurons identified as responsive to Summer in the separate single neuron analysis, we selected a subset of neurons, ensuring an even distribution across both patients and regions. Similar to the threshold model, most models exhibited minimal prediction performances (see Fig. S6). In summary, our decoding network achieved consistent and statistically significant decoding performance on the population level exceeding chance level for all labeled movie features.

### Choice of architecture

Our pipeline uses a recurrent neural network (LSTM) to process spiking data as a time series of event counts. We binned spike counts into 80 ms intervals, covering 800 ms before and after label onset, creating sequences with a total length of 20 and a dimensionality of 2286 neurons. We trained a separate model for the prediction of each label, with individually optimized hyperparameters. Given the high degree of imbalance for some labels (i.e., the character McKenzie only appears in 10 % of the movie’s frames), we oversampled the minority class during training to mitigate the effects of the uneven distribution. We additionally employed a 5-fold nested cross-validation procedure and carefully selected samples to avoid correlations between samples, as discussed in more detail in the following section. See Methods for additional details regarding model training and architecture.

We compared the LSTM’s decoding performance to a simpler logistic regression model, i.e. a linear method that does not consider the neuronal activity as a sequence of spike counts, and therefore ignores the temporal information and non-linear dynamics (Fig. 3c). Apart from this, the setup and data split for both pipelines were identical. The logistic regression model showed lower performances for the Scene and Camera Cut features (by 0.1 and 0.07, respectively), whereas no drop for character-related or location-related features was observed. To test our hypothesis that temporal information within spike sequences influences visual transition decoding, we assessed trained models using temporally-modified test data (sequence order of the spike trains was randomly shuned, see Methods for more details) (Fig. 3d). A pattern consistent with the LSTM decoding results emerged, with a noticeable decline in performance, especially for the visual transition labels.

### Avoiding spurious decoding performance by introducing temporal gaps

Since each frame of the movie shares a high degree of similarity with neighboring frames, we controlled for the temporal correlations in the annotated features induced by the continuous nature of the stimulus. We divided the dataset into training, validation, and test sets, ensuring a gap of 32 s between samples from different sets to minimize temporal correlations (split visualized in Supp. Fig. S7). We investigated the impact of these gaps by decoding the character Summer using varying gap lengths while keeping the number of samples comparable. We observed that smaller gaps result in substantially higher decoding performance on the test set, raising concerns about potential data leakage between training and test sets. For instance, a random split without any temporal gaps achieves an almost perfect score of 0.99 *±* 0.0005. However, as temporal gap sizes increase to 32 s, the performance drops precipitously to 0.31 *±* 0.06 (Fig. 3e; additional metrics in Supp. Fig. S9). This might explain the higher decoding performance for a comparable task in Zhang et al. ^24^, which did not report the use of temporal gaps for model evaluation. All subsequently reported results refer to the performance on the held-out test data using 5-fold cross-validation, with data splits incorporating the most conservative temporal gap of 32 s (see Methods). Our analysis underscores the importance of appropriate architecture selection and careful data preparation in complex datasets such as ours, as these choices can exert a significant impact on the results.

#### Patient-wise decoding performance

The neuronal population analyzed thus far has been pooled from 29 patients, yielding a total of 2286 neurons. Decoding from a pooled population, rather than from individual patients, improves network stability by aggregating activity across a larger neuronal set and enhances the signal-to-noise ratio. However, using this pooled population (or “pseudo-brain”) obscures the patient-wise contributions to decoding, which could vary due to the difference in neurons recorded per patient (units per patient range from 30 to 137) or due to differences in the semantic space of recorded neurons. To test for such differences, we assessed decoding performance on a per-patient basis, and analyzed each participant’s neuronal population to see if key decoding information is widely distributed or driven by a particular subset.

Decoding performance was obtained for three label categories—Summer, Scene Cuts, and Indoor/Outdoor—representing characters, visual transitions, and locations. To minimize computational load, we retrained a simpler logistic regression model on each patient’s neurons, and achieved performance comparable to a more complex recurrent neural network but with lower computational costs. The results are illustrated in Fig. 4. Generally, decoding performance was lower at the individual patient level compared to the pooled neuronal population, with no single patient matching the performance of the aggregated data. For the Summer label, we observed substantial variability in decoding performance across patients, with some patients showing near-zero accuracy. However, certain patients (specifically 7,10,15, and 22) achieved decoding performances exceeding 0.2, compared to the overall pooled performance of 0.28. This variability was less pronounced for the Scene Cuts and Indoor/Outdoor labels. For the Scene Cuts label, the already low pooled performance declined further in the per-patient analysis, with patients 2,10,19, and 20 showing slightly better results, while most demonstrated minimal decoding accuracy. The Indoor/Outdoor label elicited a consistently higher accuracy across patients, matching the overall higher decoding performance achieved with the pseudo-brain population. Notably, patients 2,13,20 and 25 achieved performances exceeding 0.2 (Cohen’s Kappa), compared to an overall pooled performance of 0.31, indicating robust neuronal responses in patient-specific subpopulations of neurons. Across all labels, the highest-performing patients vary, and no single patient showed consistently superior performance across all three.

**Figure 4.**
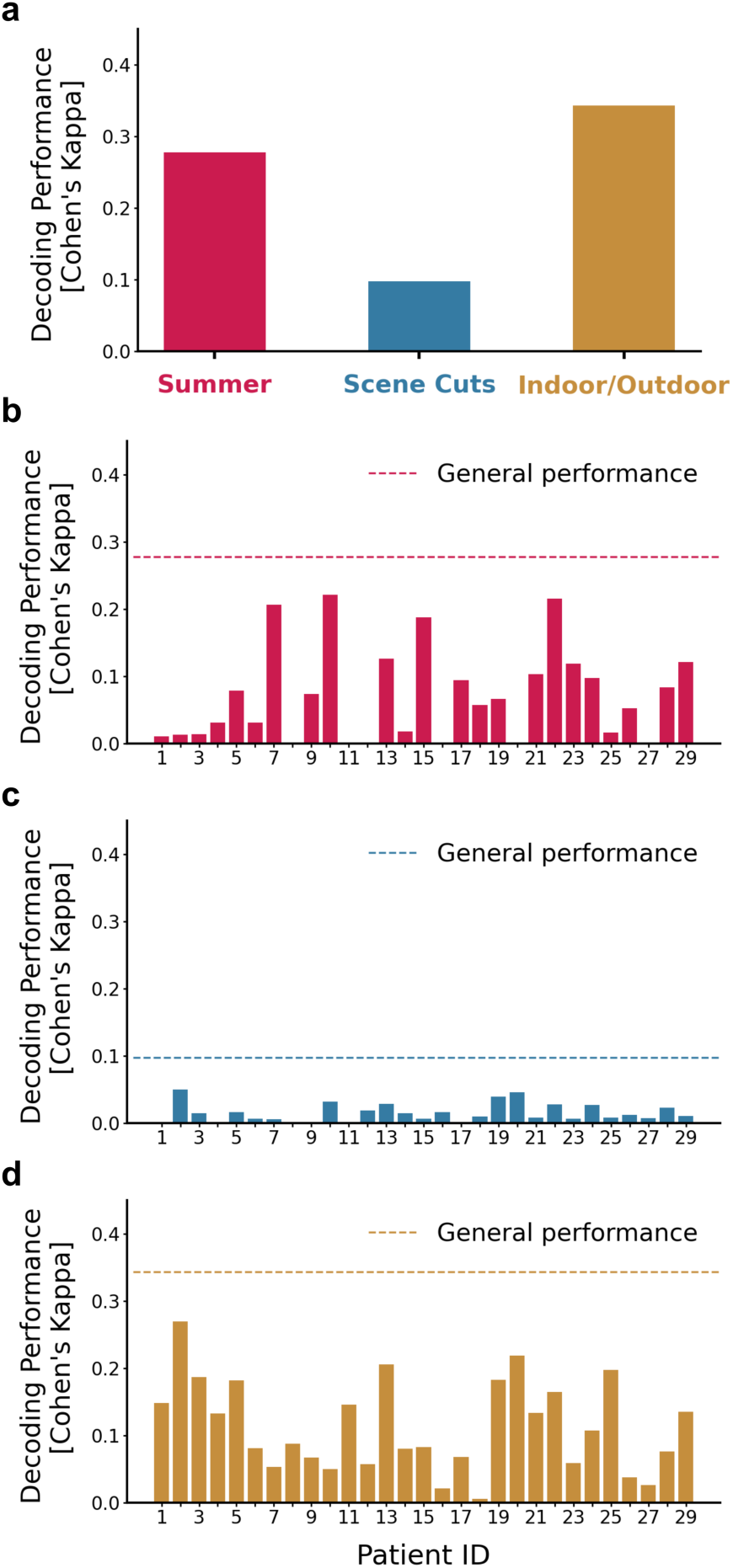
Patient-specific decoding performance Decoding performances for the main character (Summer), visual transitions (Scene Cuts), and location (Indoor/Outdoor) are reported using Cohen’s Kappa and compared to the performance obtained from the total population (pooled across all patients, dashed line). a) Decoding performance based on the total population of 2286 units, with neurons pooled across all patients. b-d) Patient-specific decoding performances for Summer, Scene Cuts and Indoor/Outdoor.

### Parahippocampal cortex drives decoding of visual transitions and location

Continuously presented stimuli offer a rich array of features. Visual transitions, such as changes in filming angle or scenery, are a commonly studied feature that demarcate the event structure of the dynamic stimulus. In movies, these transitions are relatively well-defined since they consist of identifiable changes in pixel values between frames and are known to elicit time-locked changes in neural activity in fMRI ^20^, iEEG ^22^, and single neurons ^11^. We investigated two types of frame-wise visual transitions: Scene Cuts (changes in scenery) and Camera Cuts (changes in filming angle). As Scene Cuts consists of visual transitions between locations or points in time and demarcate narrative episodes within the movie, they are related to location but not exclusively. We compared this label to a more straightforward location-related feature, Indoor/Outdoor, which indicates whether a given frame depicts an indoor environment or not. Examples of Scene versus Camera Cuts as well as Indoor/Outdoor scenes are shown in Supp. Fig. S2.

To investigate region-wise differences, we trained separate decoders for neurons in the amygdala (A), hippocampus (H), entorhinal cortex (EC), and parahippocampal cortex (PHC) of the MTL. We excluded the piriform cortex (PIC) due to its relatively lower number of recorded neurons (Fig. 1b). We observed a clear dominance of the parahippocampal cortex for both types of visual transitions. The decoding performances reached 0.21 *±* 0.02 for Scene Cuts and 0.19 *±* 0.03 for Camera Cuts, respectively, when restricting the decoding to the parahippocampal cortex as opposed to 0.20 *±* 0.03 and 0.18 *±* 0.01 when decoding from the full population. The other regions showed a lower but above-chance decoding performance. Similarly, the parahippocampal cortex yielded the highest performance for predicting indoor versus outdoor content and reached a performance of 0.30 *±* 0.06, comparable to the performance on the full population (0.31 *±* 0.06). Hippocampus was the second strongest region with a high performance of 0.26 *±* 0.04. For entorhinal cortex and amygdala, we observed lower performances of 0.15 *±* 0.05 and 0.1 *±* 0.03, respectively.

In summary, the parahippocampal cortex consistently achieved the highest decoding performance for labels associated with visual transitions and location, in line with prior research on the MTL ^11^. Given that both Scene and Camera Cuts are linked to sharp visual transitions, we anticipated that earlier processing stages in the MTL would show better decoding performances than later processing stages. However, despite the clear dominance of the parahippocampal cortex, our results show that other regions achieve lower, but nonetheless significant performance when detecting event structure and setting information.

### Amygdala drives decoding of character presence

The MTL carries information about the identity of specific individuals, in addition to general person-related categories or attribute, primarily through the tuning of individual neurons ^31–33^. Unlike visual transitions, character identities are a semantic feature which rely on both visual attributes and higher-level abstract representations. To investigate character-driven representations at the population level, we analyzed neuronal activity during the presence of the movie’s three main characters, Summer, Tom, and McKenzie, as well as the more general concept of any character appearance (Persons, see example frames in Supp. Fig. S2). While Summer’s appearance throughout the movie frames is balanced (50*/*50), the remaining labels are highly imbalanced: Tom and Persons appear in the majority of frames (80*/*20 and 95*/*5), while McKenzie is predominantly absent (10/90) (Fig. 1c). Despite the imbalances, we observed significant decoding performances for all four character labels ranging between 0.23 and 0.36 (Fig. 3c). Notably, decoding performance for character identities—despite being abstract and variable—exceeded that of visual transitions (0.20 and 0.18).

### Distribution of information across MTL regions

To investigate whether characters were primarily processed in a specific MTL region or in all regions equally, we conducted a similar analysis as before by retraining on region-specific activity (Fig. 5a). Our results show that all four tested regions carry information about the character’s identities, enabling decoding at above-chance levels (*p <* 0.001, permutation test, see Methods). The amygdala and parahippocampal cortex showed the highest decoding performances for Summer and Tom, respectively, approaching levels similar to decoding from the full population. However, the distribution of information among the other regions was less consistent and varied across labels (Fig. 5a). The hippocampus had the lowest performance for Summer (0.09 *±* 0.04) and Tom (0.05 *±* 0.05), while the entorhinal cortex performed lowest for McKenzie (0.12 *±* 0.02). For Persons, the parahippocampal cortex dominated with decoding performance comparable to the full population (0.36 *±* 0.05), while other regions ranged between 0.14 and 0.21.

**Figure 5.**
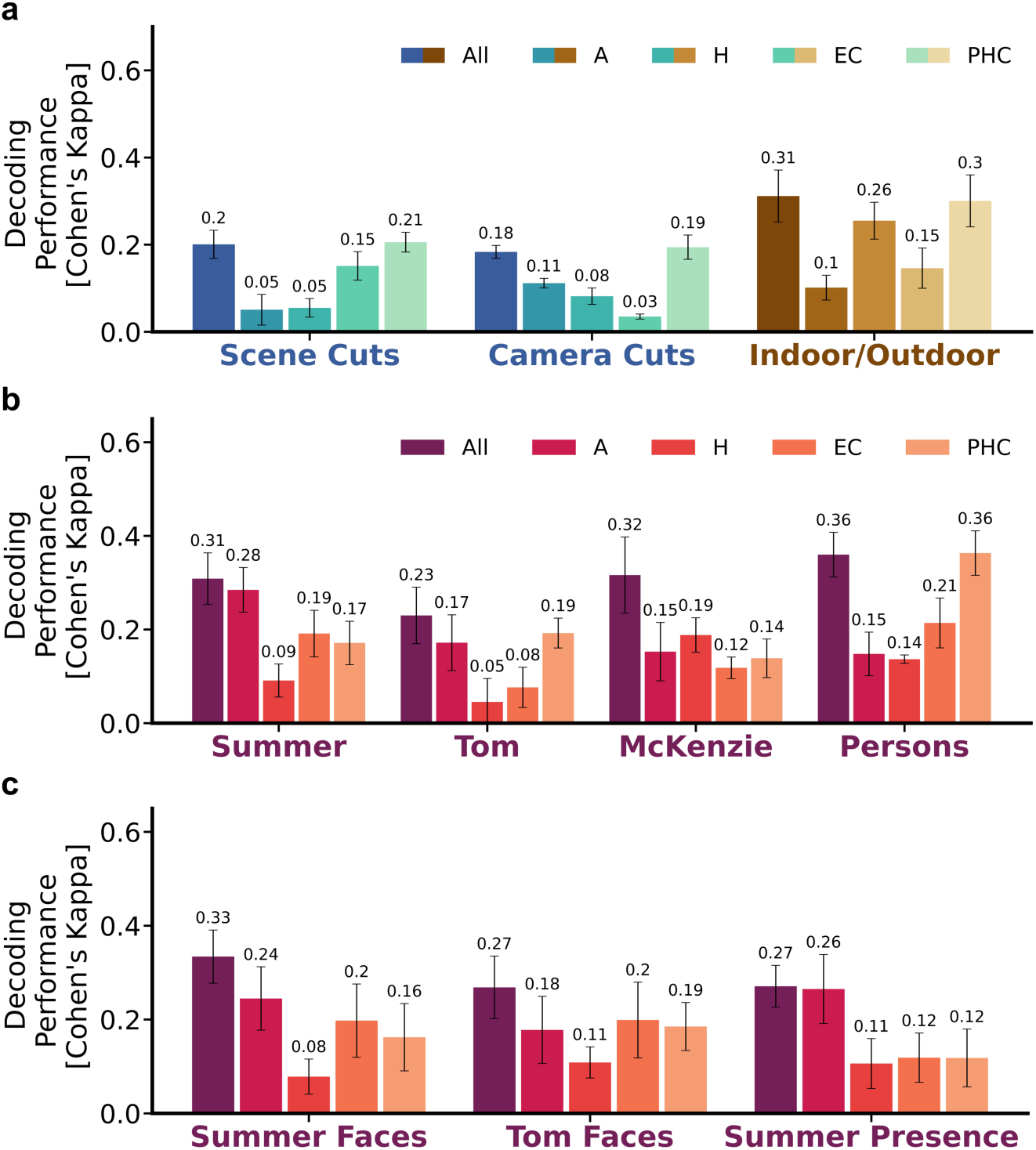
Semantic information is distributed differently across MTL regions based on category Decoding performances for semantic features, by MTL region. All performances were significantly better than chance level with an alpha level of 0.001 (reported performance using Cohen’s Kappa, mean performance across five different data splits, error bars indicate standard error of the mean. a) Decoding performances for visual transitions and location features. b) Character visibility could be decoded from the entire population of neurons, and with variable performance when training only on individual MTL regions. c) Decoding performances for face-specific character appearances and Presence features, by region.

### Differences in character’s visual appearances

Since the labels indicate the presence of a given feature within a natural scene rather than a single exemplar shown in isolation, the visual appearance of the labeled entity varied substantially during the movie.

To better control for visual appearance and examine decoding differences across various levels of character presence, we created the additional labels Summer Faces, Tom Faces, and Summer Presence (see Methods for details on annotation creation). As with the character labels, we trained a decoding network on both the full neuronal population as well as the four individual regions (Fig. 5b). Face labels for both characters elicited a slight improvement in performance compared to the full neuronal populations, with Summer increasing from 0.31 *±* 0.06 to 0.33 *±* 0.06 and Tom increasing from 0.23 *±* 0.06 to 0.27 *±* 0.07. Regionally, the distribution remained consistent with the general character labels, except for Tom Faces, where the entorhinal cortex had the highest performance. The hippocampus performed weakest for both Summer Faces (0.08 *±* 0.04) and Tom Faces (0.11 *±* 0.03). The Summer Presence label had an overall performance of 0.27 *±* 0.04, slightly lower than the general Summer label, with the amygdala clearly dominating the regional distribution (0.26 *±* 0.07). For face labels, distributions were more similar across regions than for the general character labels. The amygdala outperformed other regions in decoding the abstract presence of Summer but performed poorly for general person appearances, where the parahippocampal cortex performed best. These findings align with previous research showing that semantically-tuned cells in the human MTL can flexibly activate when their preferred conceptual category is indirectly invoked ^34^.

### Responsive neurons drive decoding of visual transitions but not decoding of characters

Although only a subset of neurons modulated firing in response to the onset of a given movie feature, we nevertheless observed significant decoding performance from the full population. This effect could result from two scenarios: a) information is distributed throughout the neuronal population and decoding does not disproportionately rely on neurons with post-onset increases in firing, or b) the subpopulation of responsive neurons informs the decoder while non-responsive neurons are ignored. We tested each scenario by dividing the full population into two corresponding subsets—non-responsive and responsive neurons—and re-training a neural network on each subset. We analyzed the labels Summer and Camera Cuts, which represent the character-related and visual transition categories, respectively. We then compared the prediction performance of each re-trained model to that of the full population to determine if the decoding of a given label took the entire population into account (scenario a) or relied on responsive neurons (scenario b) (Fig. 6). The subsequent analysis evaluates these subsets both at the full population level and within the restricted context of MTL regions.

**Figure 6.**
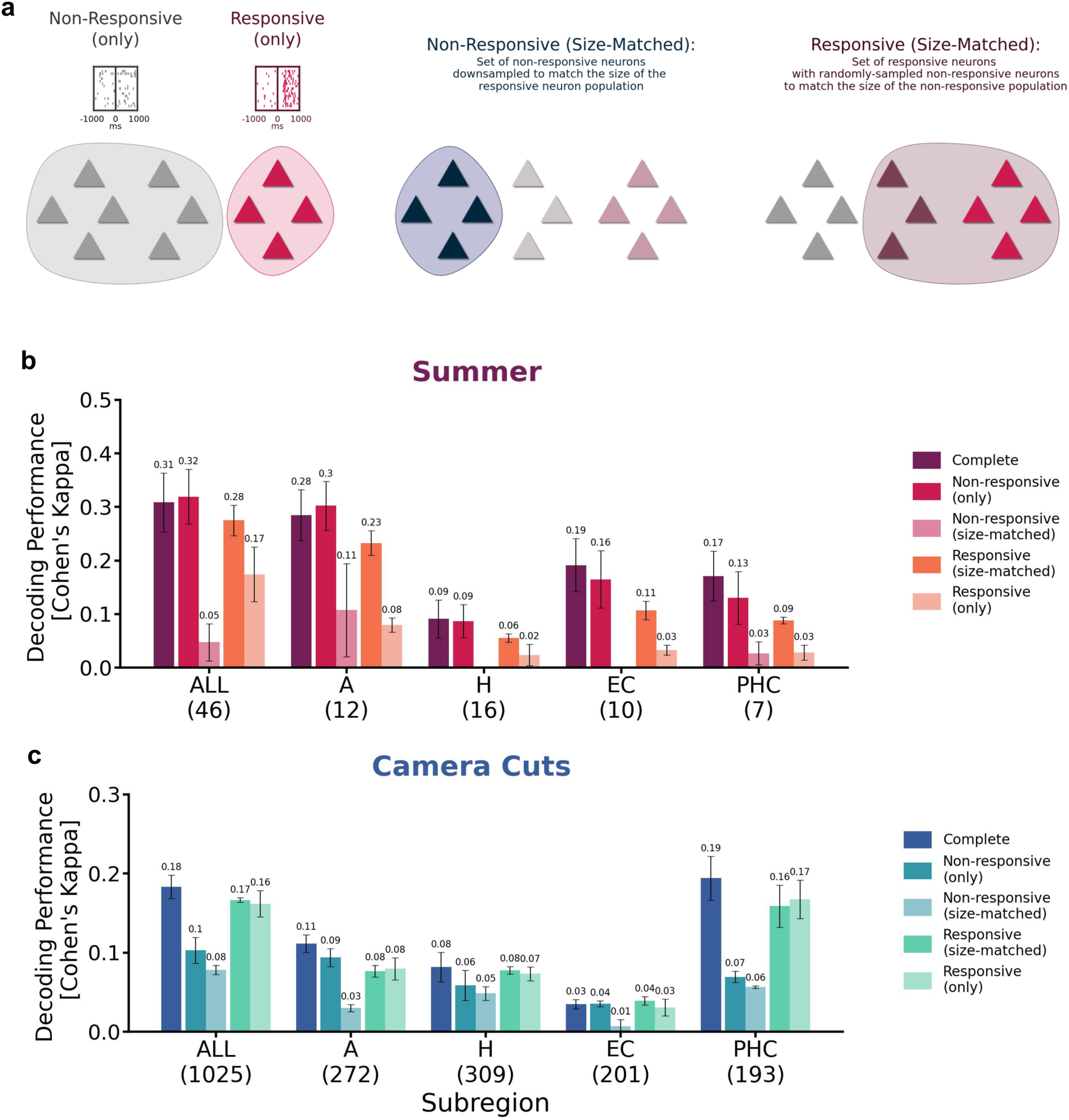
Responsive neurons drive performance for visual transitions, but not characters. To assess the contribution of the responsive neurons on decoding (identified in *Stimulus-aligned responsive neurons found primarily in parahippocampal cortex*), we compared the decoding performance for subpopulations which did or did not contain these neurons. a) Illustration of neuronal sets used in the decoding comparisons (triangles represent neurons). An example of the complete population is shown in the left-most section, which depicts *Non-responsive (only)* and *Responsive (only)* cells, with a simulated example of a respective peri-stimulus time histogram (onset raster, grey and magenta). A *Non-responsive (size-matched)* group (middle section) was randomly subsampled from the total population to have a size-matched comparison to the total set of responsive neurons. The *Responsive (size-matched)* set (right-most section) consisted of all responsive neurons padded with randomly selected non-responsive neurons to match the total *Non-responsive (only)* population. b,c) Decoding performances for discussed subpopulations for the character label Summer and Camera Cuts. Number of responsive neurons for the respective subpopulation reported in parentheses.

### Decoding with non-responsive versus responsive neurons

First, we tested prediction performance using only the non-responsive neurons to determine if similar decoding could be obtained without neurons which significantly modulated firing after the onset of a feature. For this subpopulation (*Non-responsive (only)*), a separate LSTM was retrained and tested. Despite the exclusion of responsive neurons, these subpopulations yielded performances comparable to those of the complete population (Fig. 6a, *Complete*) for the character Summer. Minimal differences in performance were observed across MTL regions, with only the entorhinal and parahippocampal cortex showing qualitative decreases. For comparison, we additionally retrained using only responsive neurons as input, and again tested the decoding performance (*Responsive (only)*). In contrast to the minimal differences observed in the *Non-responsive (only)* model, restricting to only responsive neurons produced a decrease in overall performance across the entire MTL and all regions. This general decrease suggests that the subset of individually responsive neurons is not the primary driver of the decoding performance observed in the full population model. Note that the total number of responsive neurons was less than the total non-responsive, so this effect may be influenced by an overall decrease in neuronal data. This difference in totals is directly addressed below by a size-matching procedure.

A different pattern emerged for the Camera Cuts feature, which exhibited the greatest performance drop when responsive neurons were excluded, for the entire MTL and for all subregions but the entorhinal cortex (Fig. 6b). This pattern was most pronounced for the parahippocampal cortex, indicating that responsive neurons carry valuable information for processing Camera Cuts in the movie. This hypothesis is further supported by the decoding performances obtained when restricting the decoding to only responsive neurons. Despite the restricted number of input neurons, the performance dropped marginally, and remained comparable to that of the complete population for the entire MTL, as well as for the amygdala, hippocampus, and entorhinal cortex regions.

### Size-matched neuronal populations: comparing responsive and non-responsive subsets

Since the total number of responsive neurons was lower than that of non-responsive neurons, we tested the performance using size-matched versions of both non-responsive and responsive subpopulations. To match the size of the responsive neurons, we randomly selected an equivalent number of non-responsive neurons (*Non-responsive (size-matched)*) and trained and tested a separate neural network. This process was repeated three times with different random selections, and we report the average performance. For both labels, Summer and Camera Cuts, the smaller size-matched non-responsive subpopulation showed an expected decrease in performance compared to the full non-responsive subpopulation. However, the results diverged when comparing the size-matched populations: For Summer, the size-matched non-responsive neurons performed comparably to the responsive neurons within individual MTL regions. Only for the complete population did the responsive neurons show a clear improvement. Conversely, for the Camera Cuts, restricting to only the responsive neurons improved performances, with decoding predictions of all but the amygdala surpassing those of the size-matched non-responsive set of neurons.

A similar pattern emerged for a size-matched version of the responsive neurons (*Responsive (size-matched)*), which we formed within region by padding the set of responsive neurons with randomly selected non-responsive neurons. For Summer, decoding from this subpopulation consistently showed a performance drop compared to the total set of non-responsive neurons across all tested regions. In contrast, for Camera Cuts, the size-matched subpopulation of responsive neurons achieved similar or better performance than the non-responsive neurons across all regions, with the complete population and parahippocampal region showing a clear dominance of the subpopulation containing responsive neurons. We additionally evaluated performances for the labels Tom, Scene Cuts and Indoor/Outdoor, which matched the effects found for Summer and Camera Cuts (Supp. Fig. S10).

In summary, our findings indicate that responsive neurons play distinct roles for different features. Neurons responsive to visual transitions appeared to carry information not equally present in other neurons. On the contrary, for character- and location-related labels, individual responsive neurons contributed less to decoding, and information appeared to be distributed either across the entire population or a subset of neurons distinct from the previously identified responsive neurons.

### Relevant information is carried by a smaller subpopulation of 500 neurons

We observed that subsets of neurons with stimulus-selective responses to characters did not account for the decoding performance of the same character. Previous research in sensory information processing suggests that relevant stimulus information is often encoded by only a subset of neurons within a population ^27,35,36^. To explore this further, we adopted a more data-driven approach to define neuronal subsets, ranking their importance using weights extracted from a trained logistic regression model and investigated the minimally sufficient number of neurons required for successful decoding.

### Ranking of neurons for the character label Summer

The weights of a logistic regression model can be used to assess each neuron’s importance in decoding, allowing for the creation of a ranking across all neurons (see Methods for more details). In contrast to an LSTM, logistic regression models are computationally less expensive to train and achieve comparable decoding results for all movie features, except for Scene Cuts and Camera Cuts, which exhibited reduced but above-chance decoding performance (Fig. 3c). For the character label “Summer,” we generated a neuron ranking from the trained logistic regression model and defined subsets of neurons by selecting those with the highest rankings.

The above LSTM and logistic regression models were trained on population data using 5-fold cross-validation, with alternating test sets for each of the five splits. However, this approach precludes an independent ranking of neurons across splits, as test data from one split may overlap with training data from another. To address this, we expanded the five original splits to 20, where each subgroup of four splits shared a common test set but varied in the allocation of training and validation data. Any analysis relying on a subset of neurons derived from logistic regression weights was exclusively assessed using the four splits that produced the ranking and shared held-out test data (see more details in Methods, and Supp. Fig. S8).

### Training on pre-selections of top-ranked neurons

We trained on progressively smaller subpopulations of top-ranked neurons for the character Summer (Fig. 7a) and observed an increase in performance when restricting the input activity to smaller populations (peak at 0.38 (Cohen’s Kappa) for 500 neurons, 21.9% of the total population). The decoding performance reported here for the entire population shows a slight variation from the previously reported value of 0.31 due to the modified nested cross-validation procedure. Further reduction of the population lead to a decrease in performance, yet high decoding performance persisted even in small subpopulations of neurons.

**Figure 7.**
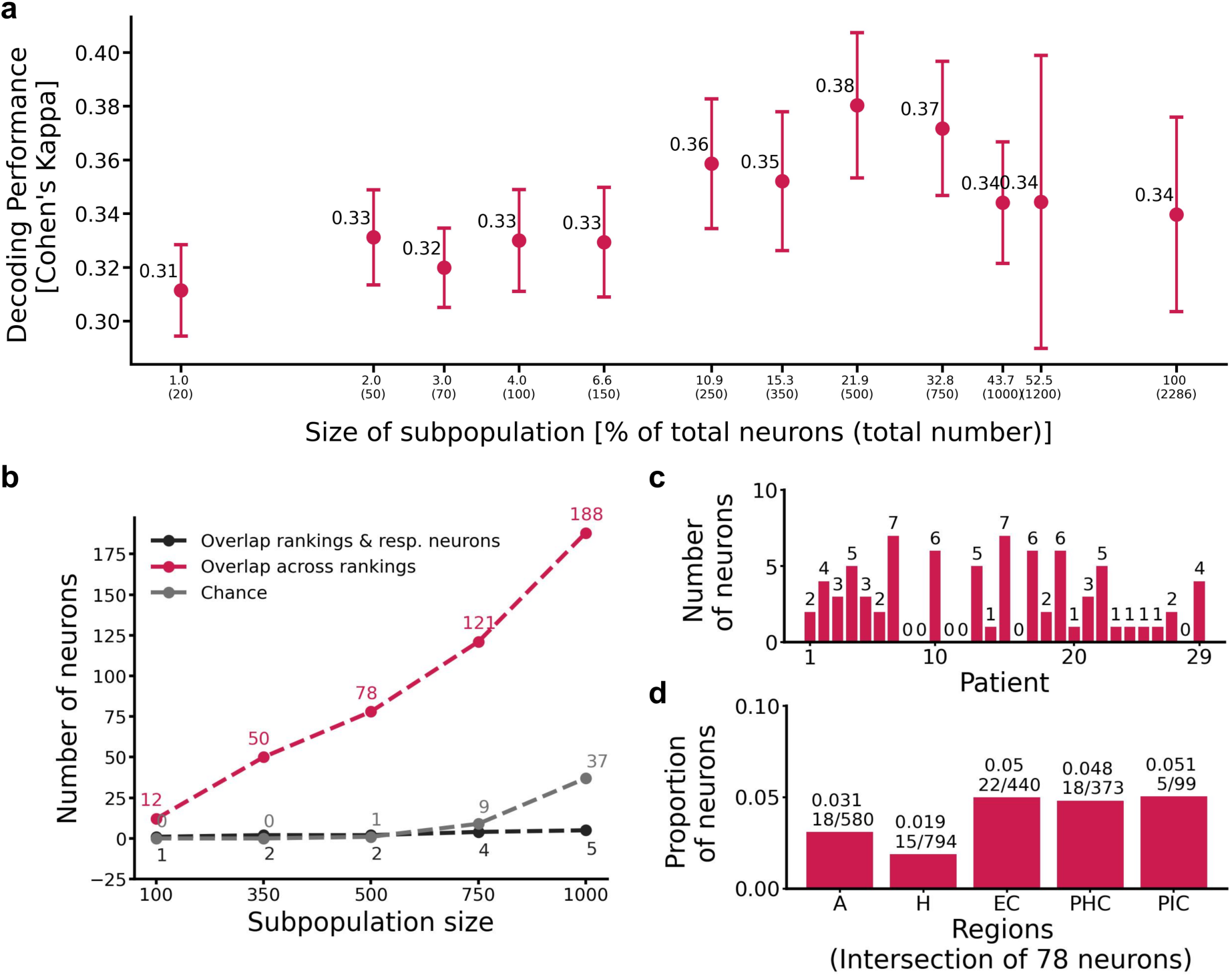
500 neurons are sufficient to reach peak decoding performance a) Decoding performances for the character Summer for subpopulations of top-performing neurons, testing various sizes ranging from 1% to 100% of the full population (absolute numbers of neurons are reported in parenthesis). Mean performance across different splits is reported, and standard error of the mean is visualized by the error bars. b) Number of overlapping neurons across rankings for different sizes of subpopulations of top-performing neurons (pink). As a baseline, we compare the number of overlapping neurons to the number expected by chance (grey), and we observed a notably higher intersection of top-ranked neurons across the splits. Additionally, the overlap between the intersection of top-ranked neurons and the previously defined responsive neurons is shown (black). c, d) Overlapping neurons (in total 78) in subpopulations of 500 top-performing neurons for each ranking were distributed across patients and MTL regions.

We observed that a ranking procedure which did not use a common test set was subject to potential cross-talk between data splits, which substantially impacted and distorted the results. Using a selection of neurons derived from non-independent training and test data led to a stark increase in performance, nearly doubling the original performance of the character Summer to 0.52 (Cohen’s Kappa) when restricting to a subpopulation of 150 top-ranked neurons. This again underscores the need to carefully prepare the data for paradigms such as ours, where data samples are highly correlated, as ignoring dependencies between training and test data can greatly skew results (see Supp. Fig. S13).

### Top-ranked neurons and their distribution across patients and MTL regions

As the selection process involved five distinct rankings of neurons, we investigated the consistency of the neuronal composition across rankings. The overlap of neurons within subpopulations of top-ranked neurons assessed across different sizes is shown in Fig. 7b. Analyzing the top-performing 500 neurons from each of the five rankings revealed a common set of 78 neurons. We compared this observed overlap to that expected by chance with random subpopulation selections (see Methods), finding a notably higher overlap. This suggests the presence of common neurons crucial for decoding the label Summer. Additionally, we also compared these selected neurons to those previously classified as responsive. The overlap between the two groups of neurons was small, suggesting that the responsive neurons defined by onset-related changes in activity may not be the primary drivers of decoding performance in general, aligning with previous observations for character labels when the decoding pipeline was restricted to responsive neurons. As restricting to 500 neurons yielded the highest decoding performance, we subsequently analyzed the resulting intersection of 78 neurons across rankings. These 78 neurons were distributed across both patients and regions (Fig. 7c,d), with no single patient or area of the MTL driving the performance. Visualizations of the spiking activity surrounding the onset of Summer for these neurons do not reveal a clear pattern of stimulus-evoked increases in firing (Supp. Fig. S12).

Our findings reveal that a core subpopulation of approximately 500 neurons drives decoding performance, while additional neurons mainly contribute redundant or noisy information. These neurons are distributed across patients and MTL regions, presenting an important avenue for future research to investigate the mechanisms underlying their organization and the specific functional roles they play in decoding processes.

## Discussion

We investigated how the human brain processes semantic and event structure in a naturalistic setting by analyzing the activity of neurons in the medial temporal lobe during the presentation of a full-length commercial movie. Although earlier work has established and characterized the role of MTL neurons in semantic representation, and the processing of dynamic stimuli via fMRI and intracranial electroencephalography, few studies have investigated how human single neurons process dynamic stimuli and none have addressed the relationship between representations on the single-neuron level and the population level.

By analyzing changes in each neuron’s activity aligned with the onset of labeled movie features, we identified groups of individual cells that adjust their firing rates in response to specific features. The most pronounced responses occurred following changes in camera angles and scenes, as these features induced activity changes in the largest number of cells. Outdoor scenes and the two main characters also elicited consistent single-unit responses, albeit in far fewer neurons. This lack of explicit single-neuron responses to characters, which might otherwise suggest selective and invariant representations such as those in concept neurons, could be explained by the study participants’ lack of familiarity with the movie. Most participants had neither seen the movie before nor encountered the actors in other media prior to its presentation as part of this study. This interpretation aligns with previous research showing that neuronal selectivity to individuals varies with familiarity and personal relevance, as photos depicting personally known individuals are more likely to elicit selective responses in MTL neurons ^37^.

We anticipated that the decoding performance for each label would reflect the pattern observed at the single-neuron level: Visual transitions would be the most accurately predicted feature, followed by setting, with character presence showing the lowest prediction accuracy. Although not every feature elicited explicit responses from a significant portion of individual neurons, the network, which takes the collective population activity as input, successfully decoded all tested features with above-chance performance. Interestingly, the decoding performance varied across features and contradicted the pattern found in the individual neurons. Despite eliciting the highest proportion of responses at the single-unit level, visual transitions showed the lowest decoding accuracy out of all tested features. Conversely, characters showed the highest decoding accuracy despite there being few individually responsive neurons. Both approaches link neuronal responses to the movie content but differ markedly in their focus, as the single-unit analysis targets the specific onset of features, emphasizing their initial activation, while the population approach processes data continuously, decoding both the onset and sustained presence of features. This methodological difference may affect the results and contribute to the observed contradiction between them. To bridge these findings, we examined how individual responsive neurons contribute to the decoding network. We hypothesized that the decoding performance for each label would be primarily driven by the subset of neurons that exhibit increased activity in response to that label. This hypothesis held true for the decoding of visual transitions, as the set of individually responsive neurons disproportionately contributed to the decoding performance when using the full population, and performance was strongly affected by their removal from the training population. In contrast, character decoding does not rely on individually responsive cells, as removing neurons that responded strongly to character onset had little impact on performance. When analyzing the network and its prediction behavior, we identified a subset of units which contributed most to the decoding of character information across models, and found that these units had little overlap with the set of neurons which increased firing after character onset. Together, these findings suggest that character-related representations relied on a population code, while visual transitions were encoded by the activity of specific neurons.

When training separately on different regions of the MTL, we observed variations in decoding performance depending on the specific content being decoded. The parahippocampal cortex achieved the highest performance in detecting visual transitions, while the amygdala performed best in predicting character-related information. These results support previous findings which identified that certain regions are more likely to respond to certain categories of images in static screenings. For example, previous studies have shown that the amygdala preferentially responds to images containing faces ^1,32^, and contains cells that selectively respond to whole faces as opposed to discrete facial features ^38^. We extended this analysis to examine different levels of character appearance, distinguishing between face visibility and the general presence of a character in a scene (regardless of face visibility). Although, the amygdala demonstrated strong performance in both cases, it most clearly drove the decoding for the general presence of a character, rather than specifically to face visibility. This effect could be due again to the unfamiliarity of the movie and its actors, as face-specific responses have been shown to form as a function of exposure ^39^.

Previous work has found that the parahippocampal cortex is especially sensitive to scene information ^40^, as opposed to objects ^40,41^, with a higher likelihood of neuronal activation when scenes feature stronger spatial layout cues, such as depth and a recognizable background ^42^. Zheng et al. (2022) identified generalized cells, termed “boundary cells”, in the parahippocampal gyrus, hippocampus, and amygdala which modulate their firing after any visual transition event. In our study, we observed significant responses to camera cuts in these same regions, which we interpret as analogous to “soft boundaries” ^11^. However, significant responses to scene cuts were only observed in the parahippocampal cortex, whereas Zheng et al. reported responses to their analogous feature (“hard boundaries”) in all measured MTL regions. In addition to their preference for scene-related images, parahippocampal neurons have been shown to respond more frequently to outdoor images compared to indoor ones ^42^. In our dataset, a small subset of parahippocampal neurons increased firing in response to the onset of outdoor scenes, whereas none showed increased activity for indoor scene onset. Despite their limited number, these responsive neurons achieved well above-chance decoding performance, albeit falling short of the performance achieved by the complete neuronal population. Further work is needed to more accurately determine if the activity of single scene-selective parahippocampal neurons in our dataset can be explained by the onset of location-related content.

Visual transitions and character content additionally differed in their use of sequence information. Through a comparison between decoding architectures, we found that temporal information in the spike trains only mattered for the prediction of visual transitions, and that ignoring or even scrambling the sequence information had little effect on character and location features. Although using temporal dynamics improved decoding performance for visual transitions, the temporal dynamics in our data, particularly those inherent to the movie stimulus, also present a significant confounding factor. Both the movie and recorded neuronal activity are subject to a high degree of correlation in time and thus require extra consideration when formulating a pipeline to ensure that the training and test data did not contain adjacent, highly correlated samples. The correlation between training and test samples artificially inflates decoding performance, and does not reflect actual generalization to unseen data. In related research by Zhang et al. (2023), where temporal distance was not considered, high decoding performances were reported on a similar task. Based on our analyses, we anticipate that their reported decoding accuracy is overestimated, and would change if sufficiently large gaps were introduced.

As our dataset consists of single-neuron activity pooled across 29 subjects, each with activity from an average of approximately 80 recorded neurons, the extent to which claims can be made about what an individual brain does is limited. In addition, the participants watched a full-length movie in an clinical setting, where neuronal activity is not solely focused on visual stimuli and likely processes additional information. Although we cannot know whether the neuronal population that we sampled is representative of the human MTL generally, it is one of the largest samples collected to date, both in terms of neurons and patients and offers a unique opportunity to understand the processing and representation of information in the MTL population activity. Despite the inherent limitations, which preclude near-perfect performances in the decoding task, the presented approach demonstrates that movie content can nevertheless be successfully decoded from such a sub-sampled population of neurons. Future work on this dataset could leverage more advanced network architectures to explicitly model between-neuron dynamics within each patient, which could better explain the gain in performance achieved moving from the single-neuron level to the population level. Additionally, in this work we focused specifically on the *visual* content of the movie. A clear next direction would be to integrate the auditory information of the movie, and possibly disentangle the contribution of visual versus audio information streams to the neuronal representation of movie features in the MTL.

## Materials and Methods

### Participants and recording

The study was approved by the Medical Institutional Review Board of the University of Bonn (accession number 095/10 for single-unit recordings in humans in general and 243/11 for the current paradigm) and adhered to the guidelines of the Declaration of Helsinki. Each patient gave informed written consent for the implantation of micro-wires and for participation in the experiment.

We recorded from 46 patients with pharmacologically intractable epilepsy (ages 19 - 62, median age 37; 26 female patients). Patients were implanted with depth electrodes ^31^ for locating the seizure onset zone for potential later resection. Micro-wire electrodes (AdTech, Racine, WI) were implanted inside the shaft of each depth electrode. Signal from the micro-wires was amplified using a Neuralynx ATLAS system (Bozeman, MT), filtered between 0.1 Hz and 9000 Hz, and sampled at a rate of 32 kHz. Spike sorting was performed via Combinato ^43^ using the default parameters and the removal of recording artifacts such as duplicated spikes and signal interference was performed via the Duplicate Event Removal package ^44^. After all data were processed, neuronal signals, experimental variables, and movie annotations were uploaded to a tailored version of Epiphyte ^45^ for analysis. Due to disruptions in the movie playback caused by clinical interruptions, 13 patient sessions were excluded from further analysis.

### Task and stimuli

Patients were shown a German dubbing of the commercial movie *500 Days of Summer* (2009) in its entirety (83 minutes). This film was chosen because the actors portraying the main characters were relatively unfamiliar to a general German audience at the time of the initial recordings. The movie was shown in an uncontrolled clinical setting, where neither gaze nor attention were directly monitored and was presented in letterbox format without subtitles on a laptop using a modified version of the open-source Linux package, *FFmpeg* ^46^, with a frame rate of 25 frames per second. Due to the length of the movie and the possibility of clinical interruptions, patients and staff were allowed to freely pause the playback. Discontinuity in playback was controlled for within Epiphyte ^45^. Pauses and skips in the movie playback were identified through the output of the modified *FFmpeg* program and used as a basis of exclusion for patients. Patients were excluded if they did not watch the entire movie, or watched the movie discontinuously.

### Movie annotations

In order to relate the content of the movie to the recorded neuronal activity, we labeled various features on a frame-by-frame basis. These labels are binary and cover the following features:

- ***Main characters:*** *Summer, Tom, McKenzie* Frames were labeled as positive if the character could be clearly distinguished by either appearance or context. Characters and persons were considered only on a visual basis (i.e., a frame in which Tom is speaking but not visible is labeled as not containing Tom).
- ***Faces:*** *Summer, Tom* Instances of a character’s face. Positive samples are frames where the character’s face is shown, while negative samples are frames where the character’s face is not visible at all. All other frames are excluded.
- ***Presence:*** *Summer* Indicates the character’s general presence in the scene, even if the character is not visible in the frame. For instance, frames are labeled as positive if the character is part of the scene but is not visible in that particular frame due to factors like the camera angle.
- ***Visual transitions:*** *Camera Cuts, Scene Cuts* Marks visual transitions in the movie. Scene Cuts correspond to changes in scenery, while Camera Cuts are primarily based on changes in the visual stimuli.
- ***Location:*** *Inside/Outside* Distinguishes between indoor and outdoor locations in the movie. Scenes that do not clearly fit into either category are excluded from the annotation.
- ***Persons*** General appearance of any person(s)

Main character, Presence, Persons, Location, and Scene Cut labels were obtained manually using the open-source annotation program *Advene* ^47^. For face labels of the characters, we developed a deep-learning pipeline for face detection and classification. As backbone we used a pre-trained neural network for face detection and feature extraction ^48^. We extended the pipeline by a classification network consisting of fully-connected layers combined with ReLU activation functions. The classification network was fine-tuned on the movie frames to classify detected faces of the main characters, including a “not known” class for faces not belonging to the main characters. In the fine-tuning process, we used the manually created character labels for the characters. Camera Cuts were labeled automatically using the open-source algorithm *PySceneDetect* ^49^, run with default parameters and manually reviewed. Camera Cuts mark a cut in the movie by labeling the first frame after cut onset as positive, resulting in cut events associated with a single frame. To adjust for the temporal latency in brain activity, cuts in the movie were associated with frames occurring within 520 ms of the cut onset. This adjustment smooths the cut labels, rendering them more comparable to what we anticipate in neuronal responses.

### Calculation of single-neuron response statistics

The “baseline” period was defined as 1000 ms prior to the onset (e.g., the entry of a character into frame) and the “stimulus” period was 1000 ms after the onset or appearance. Pseudo-trials with baseline periods containing frames depicting the label of interest were excluded. Responsive neurons were identified using a modified bin-wise signed-rank test ^8^. The spiking activity across pseudo-trials was aligned to the stimulus onset. The baseline period was binned by 100 ms, and the normalized firing rate of the baseline period was compared to nineteen overlapping 100 ms bins defined across the stimulus period using a Wilcoxon signed-rank test (alpha = 0.01, SciPy wilcoxon ^50^, Simes corrected ^51^). Additionally, a neuron was required to have spiked during at least one third of the total pseudo-trials to be tested (otherwise, assigned *p* = 1).

### Cluster permutation test

A cluster permutation test ^26^ was used to test the difference in firing rates between the responsive and non-responsive subsets of neurons. Using the calculated response statistics, neurons were divided into two conditions: responsive (*p ↑* 0.01) and non-responsive (*p >* 0.01) for a given label. Activity for a neuron was averaged across bins, yielding a single vector of mean spike counts (spikes / 100 ms) spanning both baseline and stimulus periods for each neuron. This vector was then z-scored relative to its own mean and standard deviation. Mean spike count vectors were combined across conditions, yielding two datasets: *A_resp_* and *A_non_*, matrices containing the summarized, binned activity for all responsive and non-responsive neurons, respectively. A bin-wise comparison between *A_resp_* and *A_non_* was performed using a two-sided t-test for independent samples (ttest_ind, SciPy), producing a t-stat and p-value for each bin. Clusters were defined as temporally adjacent bins with *p ↑* 0.005 and t-stats were summed within clusters. The procedure was adapted to allow testing of multiple clusters, so no clusters were excluded at this stage. One set of 1000 permutations were performed by randomly assigning neurons to *A_resp_* and *_Anon_* such that the total number of neurons in each group was conserved and the bin-wise testing procedure was performed on each permuted dataset. Each cluster was compared to the resulting histogram of permuted t-stats. P-values for each cluster were defined as the number of permutations with a higher summed t-stat relative to the total number of permutations, and a p-value less than 0.05 was considered significant.

### Decoding from population responses

For decoding, we used population responses (input) to predict the corresponding concept labels (output). We excluded the credits from the dataset and focused solely on the narrative content. The movie was presented at a frame rate of 25 frames per second, with each frame lasting 40 ms. In total, 125, 743 frames were shown. The spiking activity of the recorded neurons was binned using a bin size of 80 ms, corresponding to the activity of two consecutive frames. Each bin was then labeled based on the first frame within that interval.

We sampled activity of the neurons before and after the onset of each frame and found that using a window of 800 ms before and 800 ms after onset yielded the best decoding performance. This resulted in data samples comprising a binary label for the concept and a spike train of length 20 (10 bins before and after the onset of the frame). Given the full population of 2286 recorded neurons, each input data sample had a dimensionality of 2286.

### Architecture

To decode concepts from the sequence of neuronal activity, we used a long short-term memory (LSTM) network ^30^, which is well-suited to process the dynamical structure of the dataset. The output of the LSTM was fed into into several fully-connected layers with ReLU activation functions to obtain binary label predictions. We found that pre-processing the raw spiking data with a linear layer of same size as the input, combined with a batch normalization layer, improved performance. We used the binary cross-entropy loss function, and optimized our network using Adam optimizer ^52^ in Pytorch ^53^ with default settings (first and second order moment equal to 0.9 and 0.999, respectively). We obtained the best results with a 2-layer LSTM, with hidden size of 32. We adapted other hyperparameters, e.g., number of linear layers, hidden sizes, batch size, learning rate, weight decay, and dropout rate, for each label.

We trained each network for 700 epochs and used the validation set to estimate the model’s ability to generalize to unseen data. As is common practice, we selected the model with the best performance on the validation set and used this for evaluation on the unseen test data. Some labels were highly imbalanced and models trained on these labels were biased towards predicting the majority class. To ensure unbiased predictions and to optimize performance, it is common to force the batches of data samples presented in the training process to be balanced by oversampling the minority class. This oversampling technique was applied to all imbalanced labels, comprising all labels except the label for the main character Summer.

### Data split

To ensure that the decoding performance was not affected by correlations between data samples, we carefully split the dataset into training, validation, and test sets. We used 5-fold nested cross-validation, and assigned 70% of the data to training, 15% to validation, and 15% to testing in each split. To avoid correlations between samples, we assigned samples from consecutive segments of the movie to each set (train/val/test) and excluded 32 seconds of data between each set (see Fig. S7). The choice of excluding 32 s was based on the results for the main character Summer (Fig. 3c). The total dataset contained 45, 200 samples, resulting in sets of 30, 800/7, 200/7, 200 samples for training/validation/test. We trained a network using the training data, optimized its hyperparameters using the validation set, and evaluated its performance on the corresponding test set. We report the final decoding performance as the average performance on all five test sets.

### Evaluation metrics

For each semantic feature, we compared the model’s prediction against the binary class labels. While accuracy is a simple evaluation metric, it is not suitable in our case due to the highly imbalanced distribution of most labels, making it challenging to compare accuracy metrics across different labels. We report all decoding performances using the Cohen’s Kappa metric ^54^ which measures the agreement between the ground-truth labels and the predictions of the network, where performance equal to zero is interpreted as chance-level and performance equal to one is interpreted as complete agreement. Cohen’s Kappa is defined as

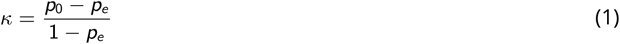

where *p*_0_ defines the relative observed agreement and *p_e_* the hypothetical probability of chance agreement among prediction and labels. The Cohen’s Kappa metric can take negative values, too, implying that the predictions are worse than chance level. The common chance performance equal to zero makes the Cohen’s Kappa a useful metric to compare performances across all labels. Additionally, we report the F1 Score, Area under the Precision Recall Curve (PR-AUC) and Area Under the Receiver Operating Characteristic (AUROC) metric in the Supplementary for all experiments. We briefly explain these metrics, including potential advantages and disadvantages for our analysis:

### F1 Score

F1 Score is a metric combining precision and recall by calculating the harmonic mean between these two. Precision and recall are determined based on a classification threshold of 0.5. Designed to perform well on imbalanced datasets, the F1 Score is particularly useful for evaluating our decoding tasks. However, the baseline for chance performance with this metric is not consistent and varies depending on the label distribution.

### PR-AUC

The PR-AUC metric, representing the area under the precision-recall curve, extends the F1 Score by evaluating performance across different threshold settings. Similarly to the F1 Score, it can be used for imbalanced datasets. However, as for the F1 Score, the PR-AUC metric is sensitive to changes in the class distributions, resulting in varying chance performance across labels. Being a reliable performance metric in general for our various classification problems, one has to bear in mind that a comparison of performances across concepts can be misleading due to the different chance baselines.

### AUROC

The Receiver Operating Characteristics (ROC) curve represents the trade-off between the true positive rate and false positive rate for various threshold settings. AUROC as the area under the ROC curve is a performance measure that is used in settings where one equally cares about positive and negative classes. Performances range in [0, 1] and it is insensitive to changes in the class distribution, which means chance performance is given by a value of 0.5. However, the metric is generally not used for highly imbalanced classification problems and an evaluation of specific labels in our analysis such as McKenzie (class distribution is 90*/*10) should be taken with caution.

### Single-neuron decoding performance

For a fair assessment of decoding from population responses compared to single-neuron activity, we evaluated the decoding performance of individual neurons under a setup comparable to that of the LSTM-based decoding network. Instead of relying on full-population responses, we established a performance based solely on the firing rates of each single neuron. Employing the same data splits, cross-validation approach, and binning procedure used for the LSTM network, we summed the binned firing rates of a neuron surrounding frame onset for the 1600 ms time window utilized by the decoding network. We then individually selected the best threshold for each neuron’s activity on the validation set, which was subsequently used to evaluate the neuron’s performance on the hold-out test set. We reported the final performance as the average of the five results obtained from the 5-fold cross-validation procedure.

### Permutation tests for decoding

To determine if the reported decoding performances were significantly better than chance, we performed two sets of permutation tests—first, we randomly shuned the labels of the held-out test set (Test Set Shune), and second, we shifted the labels while preserving the order of the test labels (Circular Shift). For both tests, the input to the decoding network remained unchanged from the non-permuted version. The only modifications made were to the corresponding feature label in the test set, which were changed in the ways outlined above, and then compared to the original network’s prediction scores. The dynamic nature of the visual stimuli implies not only a strong correlation within the neuronal activity but also suggests a temporal correlation for the feature labels. By modifying only the labels, we ensured that the temporal information embedded in the neuronal data remained unaffected by the permutation test. In the following, we test the significance of both scenarios: one where the temporal correlation within the labels is disrupted (Test Set Shune), and another where the temporal correlation within the labels is maintained (Circular Shift).

### Test set shune

The first type of permutation, and the one reported in the main text, consisted of randomly shuning labels in the test set and evaluating the predictions of the models on those. We compared the performance of the model on the heldout test set to a null distribution generated by evaluating the model on the test set with shuned labels (*N* = 1000). We calculated the probability of our observed performance under the null distribution to obtain the p-value:

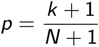

where *k* is the number of performances on permuted data outperforming the ground-truth performance of the model. This p-value provided the basis of comparison for describing significant decoding results in our main analyses. By preserving the temporal structure during the model’s inference process and only disrupting temporal correlations within the concept labels used for evaluating the model’s predictions, we consider this assessment of significance to be the most appropriate for our data setup.

### Circular shift

The second type of permutation was performed as a comparison for the above *Test Set Shufle*, as it is a standard method in human single-neuron studies (see ^55, 56, 44^). Circular shift permutations maintain both the temporal relationship of neuronal data as well as that of the stimulus information, here the concept labels. Rather than randomly shuning the labels in the test set, we applied a circular shift of labels that maintained the intrinsic temporal structure. The shifting size was randomly chosen *N* = 1000 times to obtain a null distribution akin to the previous test. This distribution was then utilized to compute the probability of our ground-truth performance and derive the p-value for assessing significance. For studies involving static stimulus presentation, wherein the stimulus is largely uncorrelated with itself, this test provides a useful way to disentangle stimulus-related effects from those endemic to the time-series information. For a comparison between the permutation results, see Supplementary Materials, Fig. S11.

### Impact of temporal information on the decoding

To assess the significance of temporal information in neuronal activity sequences, we evaluated the trained models using temporally-altered test data. The input to the decoding network included neuronal activity from 800 ms before to 800 ms after the onset of the frame, divided into 20 bins. For the temporally-altered test data, we randomly shuned the sequence order of the 20 bins (applying a consistent permutation for all neurons and data samples) and evaluated the pre-trained models on the modified test data. This procedure was repeated 100 times, and the performance was averaged. The final performance was calculated by averaging the results across the five data splits, using the standard error of the mean (SEM) as the measure of variance.

### Logistic regression models and evaluation of neuron contribution to decoding

We compared the decoding performance of the LSTM to that of a logistic regression model. The dataset used to train the logistic regression model was identical, barring one key change: the spike trains provided to the LSTM, initially of dimension (20, 2286), were reduced to a single bin representing the total number of spikes around a frame. The data then had a revised shape of (1, 2286), and no longer incorporated temporal information. We trained a logistic regression model using the liblinear solver implementation in Scikit-learn ^57^. For training, we employed an L1 penalty and z-scored the neuronal activity per neuron using the mean and standard deviation of the training data. We utilized a nested 5-fold cross-validation, and separately optimized the regularization strength for each data split. The final decoding performance was calculated by averaging the performance across all five test sets.

### Logistic regression weights for evaluating individual neuron’s contribution

Applying an L1 penalty during training enforced feature sparsity, which facilitated the interpretation of input feature importance through the model’s coefficients. A ranking of neurons was generated by evaluating the coefficients of the trained logistic regression models. Caution is needed when combining logistic regression weights from models trained on different splits due to the alternating test sets in each split, which sometimes include data used for training in another split’s test data (see Fig. S7). To avoid any interference between training and test data across splits while still accounting for the temporal variation in our data, we further modified the splits used for cross-validation. The original five splits were extended to 20 splits in the following way: each of the original five splits was further divided into four sub-splits which shared a common test set but alternated the division of training and validation data (see visualization in Fig. S8). Any analysis based on neuron selection using logistic regression weights was evaluated exclusively on the corresponding four splits that generated the ranking and shared held-out test data. Final decoding performances for such analyses were derived through a nested cross-validation procedure. This involved initially averaging the decoding performances of models that shared a common test set (i.e. averaging across each set of four sub-splits) and then averaging the resulting five performances. In short, our training procedure for the logistic regression analysis consisted of training 5 *↓* 4 = 20 splits, and thus 20 models, where each group of 4 sub-splits shared a common test set.

Neurons were ranked according to their logistic regression coefficients across each set of four sub-splits. Given that there are a total of five such groups, this lead to five distinct rankings. To obtain each ranking, we combined the coefficients of the four trained models using a two-step procedure:

1. Partition the neurons into separate subsets, based on the number of models for which a neuron had a non-zero coefficient (e.g. one group of neurons which had non-zero coefficients on all four sub-splits, then all three sub-splits, etc.).
2. Within each subset of four models, use the average of the absolute coefficients across the five splits to obtain a subset-specific ranking.

By concatenating the partitions of ranked neurons, we obtained a comprehensive ranking of all neurons. The neuron ranked highest displayed non-zero coefficients in all four models (corresponding to four sub-splits) and possessed the greatest average absolute coefficient value among neurons activated in all four models.

### Intersection of top-performing neurons and chance-level overlap

In our analysis, we restricted the decoding to subpopulations of neurons that were derived through a ranking of weights of a trained logistic regression model. To ensure that the selection of neurons was independent of the test data, the neuron selection procedure was based on five rankings derived from distinct data splits, each paired with fixed test data. We evaluated the intersection of neurons across subsets of top-ranked neurons from the five rankings, evaluated for varying subpopulation sizes (Fig. 7b). For instance, a comparison of the top-performing 500 neurons from each of the five rankings revealed a set of 78 common neurons.

As a reference point for comparison, we report the average count of overlapping neurons anticipated when randomly selecting sets of 500 neurons five times from the entire population (denoted as chance level of overlapping neurons in Fig. 7). In the previously mentioned scenario involving subpopulation sizes of 500, the expected number of overlapping neurons is equivalent to one. Mathematically, this is computed as follows: the full population consists of *N* = 2286 neurons. We refer to the size of the subpopulation as *k* = 500 and the number of total rankings *m* = 5. For each neuron *n_i_* in the subpopulation, we define a random variable *X_i_* _,500_ as follows:

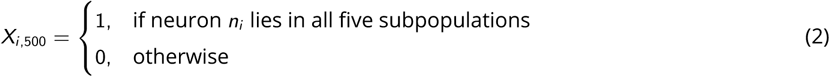

We observe that 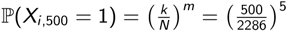. The expected value of *X_i,_*_500_ is given by

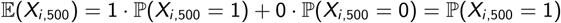

We define 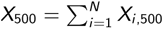 as the number of overlapping neurons across all five rankings. Since the random variables are independently and identically distributed, this implies

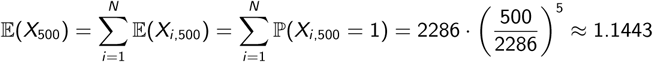

Analogous calculations for *k* = 100, 350, 750, 1000 yield

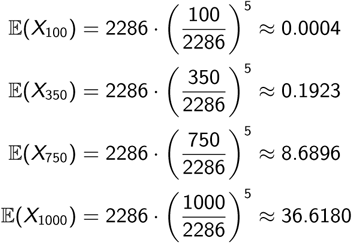

Thus, we derive the chance baselines as 0, 0, 1, 9, and 37 for subpopulation sizes of 100, 350, 500, 750, and 1000, respectively.

### Decoding on regions of the MTL

We compared the decoding performance when using the activity of all 2286 recorded neurons to the performance when only using activity from specific regions of the MTL. These regions are the amygdala (580 neurons), hippocampus (794), enthorinal cortex (440), and parahippocampal cortex (373). To use activity from a particular region, we limited ourselves to the activity of neurons in that region and reduced the input dimension to match the number of neurons in the region. The network architecture and data splits remained the same as when using activity from the full population, but the hyperparameters were optimized for the reduced dataset and given label. Training, validation, and test set sizes remained the same as the full dataset condition. In summary, decoding from different regions differed from full population decoding primarily due to reduced input data dimensionality: from a spike train of length 20 and dimension 2286 to a spike train of the same length but with a dimension reduced according to the number of neurons in the specific region.

## Acknowledgments

We would like to thank Tamara Müller and Aleksandar Levic for their contributions to the data analysis framework. This work was supported by the German Federal Ministry of Education and Research (DeepHumanVision, FKZ: 031L0197A-C; Tübingen AI Center, FKZ: 01IS18039), as well as the German Research Foundation (DFG) through Germany’s Excellence Strategy (Cluster of Excellence Machine Learning for Science, EXC-Number 2064/1, PN 390727645) and SFB1233 (PN 276693517), SFB 1089 (PN 227953431), SPP 2411 (PN: 520287829), and MO 930/4-2.

## Data Availability Statement

The data will be made fully available in a separate publication following the release of this manuscript. Code will be made available on GitHub.

## Author Contributions

### Conceptualization

Franziska Gerken, Alana Darcher, Pedro J Gonçalves, Rachel Rapp, Ismail Elezi, Johannes Niediek, Stefanie Liebe, Jakob H Macke, Florian Mormann, Laura Leal-Taixé

### Data acquisition

Alana Darcher, Johannes Niediek, Thomas P Reber, Marcel S Kehl, Stefanie Liebe, Florian Mormann

### Data curation

Alana Darcher, Marcel S Kehl

### Methodology

Franziska Gerken, Alana Darcher, Pedro J Gonçalves, Rachel Rapp, Ismail Elezi, Stefanie Liebe, Jakob H Macke, Florian Mormann, Laura Leal-Taixé

### Formal analysis

Franziska Gerken, Alana Darcher

### Funding

Jakob H Macke, Florian Mormann, Laura Leal-Taixé

### Software

Franziska Gerken, Alana Darcher, Johannes Niediek, Thomas P Reber

### Project administration

Jakob H Macke, Florian Mormann, Laura Leal-Taixé

### Supervision

Pedro J Gonçalves, Ismail Elezi, Stefanie Liebe, Jakob H Macke, Florian Mormann, Laura Leal-Taixé

### Writing - original draft

Franziska Gerken, Alana Darcher

### Writing - review and editing

Pedro J Gonçalves, Rachel Rapp, Ismail Elezi, Stefanie Liebe, Jakob H Macke, Florian Mormann, Laura Leal-Taixé

