## Supplementary for "Decoding movie content from neuronal population activity in the human medial temporal lobe"

### Supplementary Materials

#### Dataset

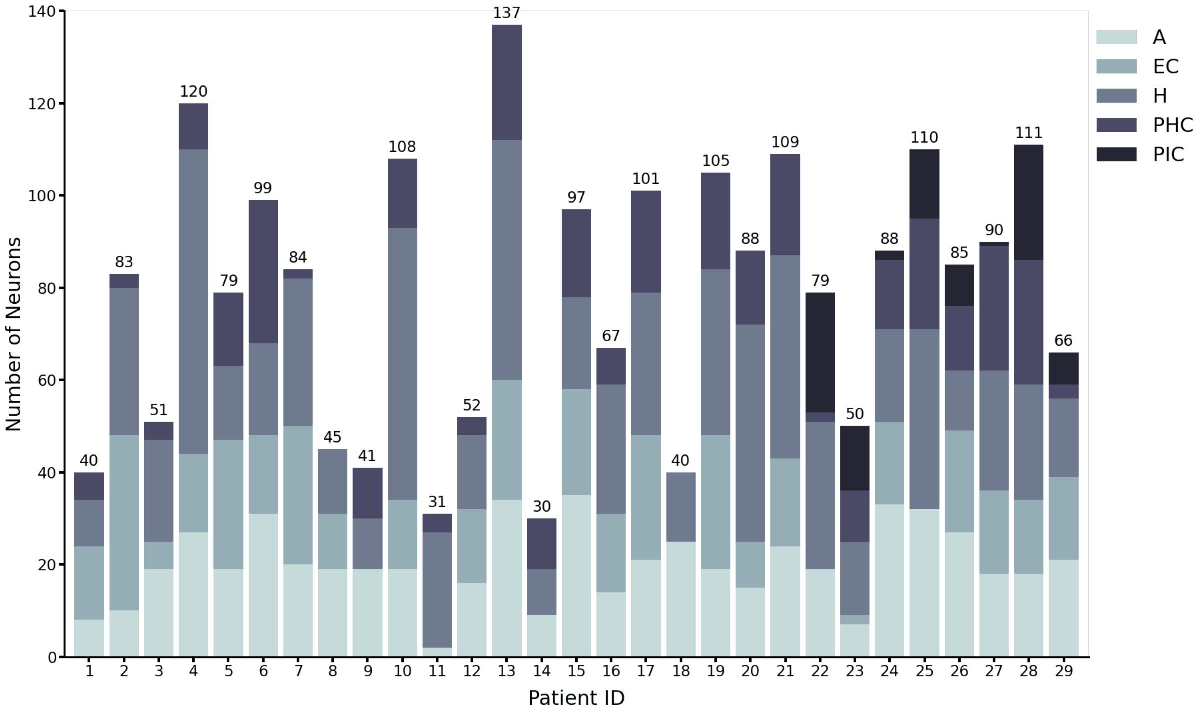


**Figure S1. Distribution of neurons across patients and by region.** Each bar shows the total number of neurons for each patient included in the dataset, with the total given at the top of the bar. The proportion of neurons in each medial temporal region for a given patient is illustrated by the size of the corresponding colored section (amygdala [A], entorhinal cortex [EC], hippocampus [H], parahippocampal cortex [PHC], piriform cortex [PIC]).

#### Annotated Features

We annotated a subset of the concepts appearing in the movie on a frame-by-frame basis, focusing on the presence of the three main characters, overall characteristics of individuals, visual transitions and indoor/outdoor distinctions. Refer to Supp. Fig. S2 for visualizations of the labeled content.

- *Visibility*
  For the general character label, visibility in any form was labeled as positive sample (Visibility row).
- *Faces*
  For face labels of a given character, any detectable appearance of their face (in which identity was contextualized by surrounding frames) was labeled as positive. The example frames show positive labeled frames for the main character Summer. Faces can appear in different sizes (Faces row).
- *Presence*
  The presence label for a given character indicates the general presence in the scene, independent of visibility in the frame. All shown frames belong to a scene where the main character Summer is present and hence labeled as positive samples. Note that due to a change of the camera angle, the character is however not visible in the middle frame (Presence row).
- *Persons*
  The more general label Persons indicates the overall visual representation of individuals in the scene, irrespective of their specific identities. Both negative and positive labeled frames are shown in the Persons row.
- *Visual transitions*
  Visual transitions occur frequently throughout the film. We differentiate between Camera Cuts and Scene Cuts:

Camera Cuts rely on a sudden frame-wise change of the filming angle or perspective and is not driven by content. They occur the most often, with an average of 1 Camera Cut every 3 seconds. Scene Cuts form a subset of the Camera Cuts and are defined by a change of scenery, i.e., location, time, environment or constellation of persons. Example frames for both transition types are shown in the Camera and Scene Cuts row, respectively.

- *Location*

  In terms of location information, we introduce a label to indicate whether a scene is indoor or outdoor. Frames that do not clearly fall into either category are marked and excluded from the analysis.

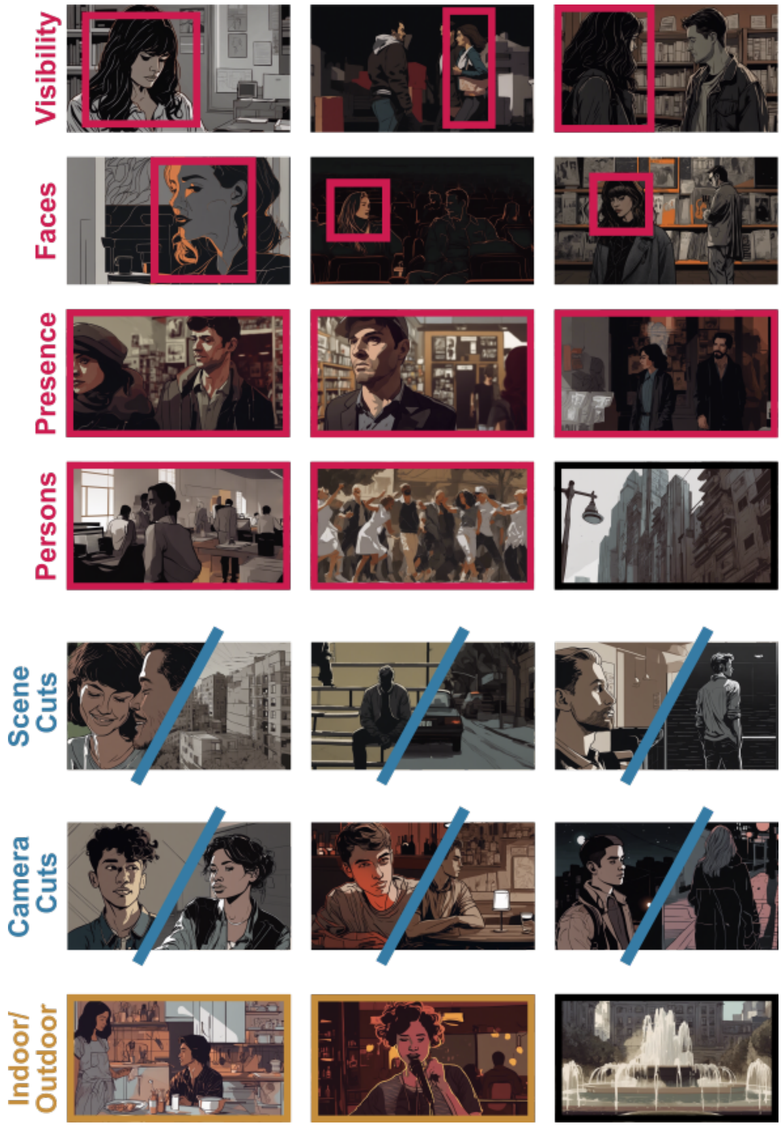


**Figure S2. Example frames of annotated movie content.** The frames capture natural scenes, with each frame encompassing a multitude of labels. To annotate the movie content, we assigned each frame a binary label denoting the presence or absence of the labeled entity. We showcase example frames and, whenever possible, indicate the decisive part of the scene using bounding boxes for positive instances of these concepts. Alternatively, a colored frame shows a scene with a positive label for a label, while those lacking the labeled content are represented in black. The movie frames used in the experiment can not be displayed due to copyright and have been replaced with images generated using stable diffusion[25].

#### Populations of single-neuron activity surrounding annotated features

By aligning the firing activity of neurons to the onset of a specific label throughout the movie, we identified responsive neurons. In our analysis, we use subpopulations of these identified neurons for the labels Summer, Tom, Scene Cuts, Camera Cuts, and Indoor/Outdoor. In Table S1 we show the respective number of those responsive neurons with additional information of their distribution across regions.

**Table S1. Proportion of responsive neurons by region and label.** The number of responsive neurons, as determined by comparing the bin-wise firing rates between baseline and stimulus periods of the pseudo-trial activity, is given in each cell. The percentage of responsive neurons for a given label and region combination is given in parentheses. The total number of neurons in a given area is given under the region abbreviation. Significant differences between responsive and non-responsive neurons, as determined by cluster permutation testing, is indicated by color. The degree of color saturation corresponds to the p-value for the largest difference in firing rate between responsive and non-responsive neurons for the label and region combination (bright green: $p \leq 0.001$, faded green: $p \leq 0.01$, grey: $p \leq 0.05$). Italics indicate label-region combinations in which the difference in curves occurred before the onset of the label.

| | Total | A (580) | EC (440) | H (794) | PHC (373) |
|---|---|---|---|---|---|
| Tom | 216 | 63 (10.86) | 39 (8.86) | 74 (9.32) | 32 (8.58) |
| Summer | 46 | 12 (2.07) | 10 (2.27) | 16 (2.02) | 7 (1.88) |
| McKenzie | 0 | 0 | 0 | 0 | 0 |
| Persons | 6 | 0 | 1 (0.23) | 1 (0.13) | 4 (1.07) |
| Tom Faces | 843 | 207 (35.69) | 174 (39.55) | 271 (34.31) | 150 (40.21) |
| Summer Faces | 634 | 160 (27.59) | 132 (30.00) | 201 (25.31) | 112 (30.03) |
| Summer Presence | 0 | 0 | 0 | 0 | 0 |
| Camera Cuts | 1025 | 272 (46.90) | 201 (45.68) | *309 (38.92)* | 193 (51.74) |
| Scene Cuts | 106 | 23 (3.97) | 18 (4.09) | 36 (4.53) | 25 (6.70) |
| Indoor | 0 | 0 | 0 | 0 | 0 |
| Outdoor | 4 | 0 | 0 | 0 | 4 (1.07) |

**Figure S3.** Single unit activity surrounding appearances of character-related labels during the movie presentation, for labels **a)** Tom, **b)** Summer, **c)** McKenzie, and **d)** Persons.

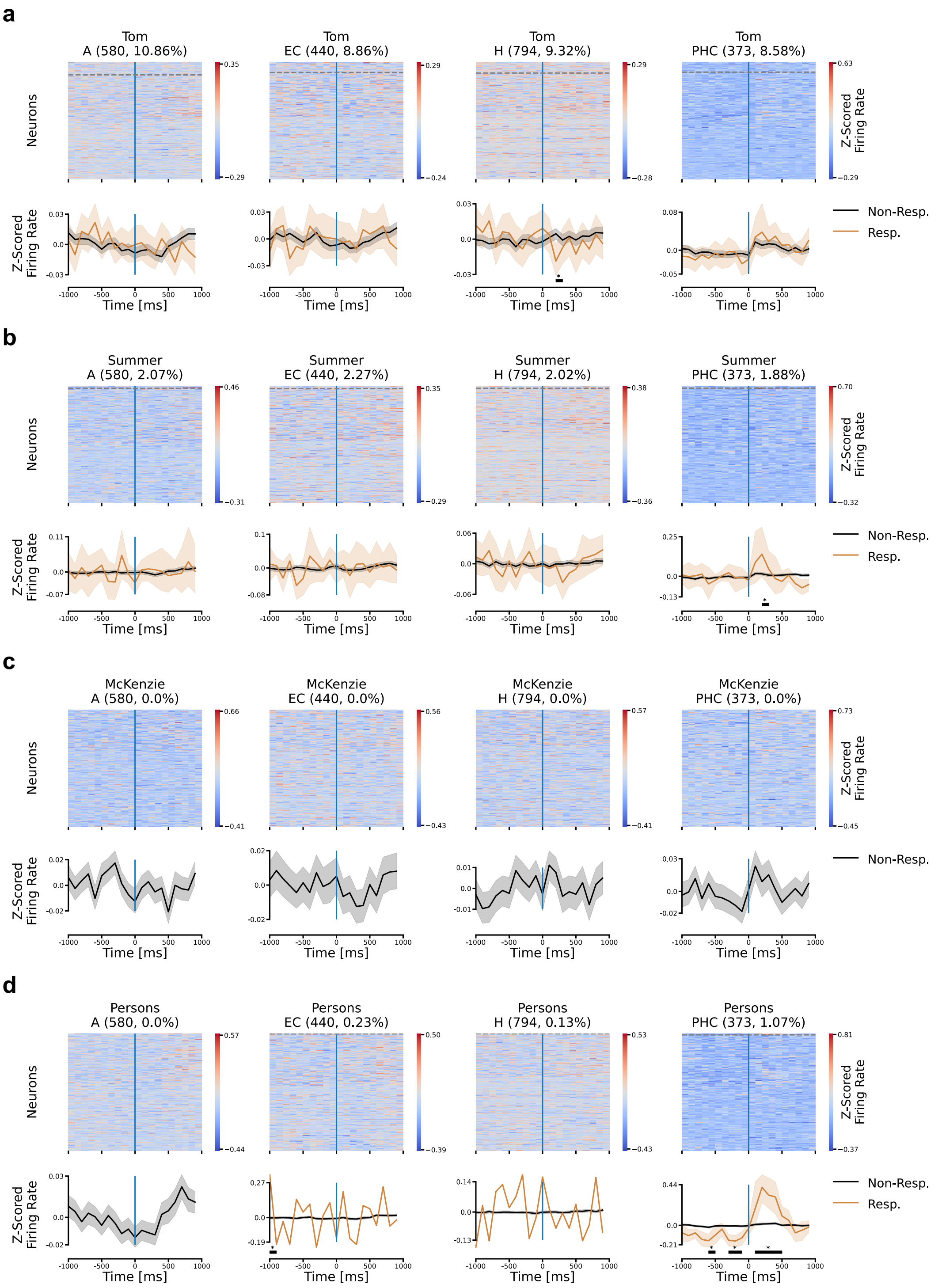

**Figure S4.** Single unit activity surrounding appearances of character attribute labels during the movie presentation. **a)** Tom Faces, **b)** Summer Faces, and **c)** Summer Presence.

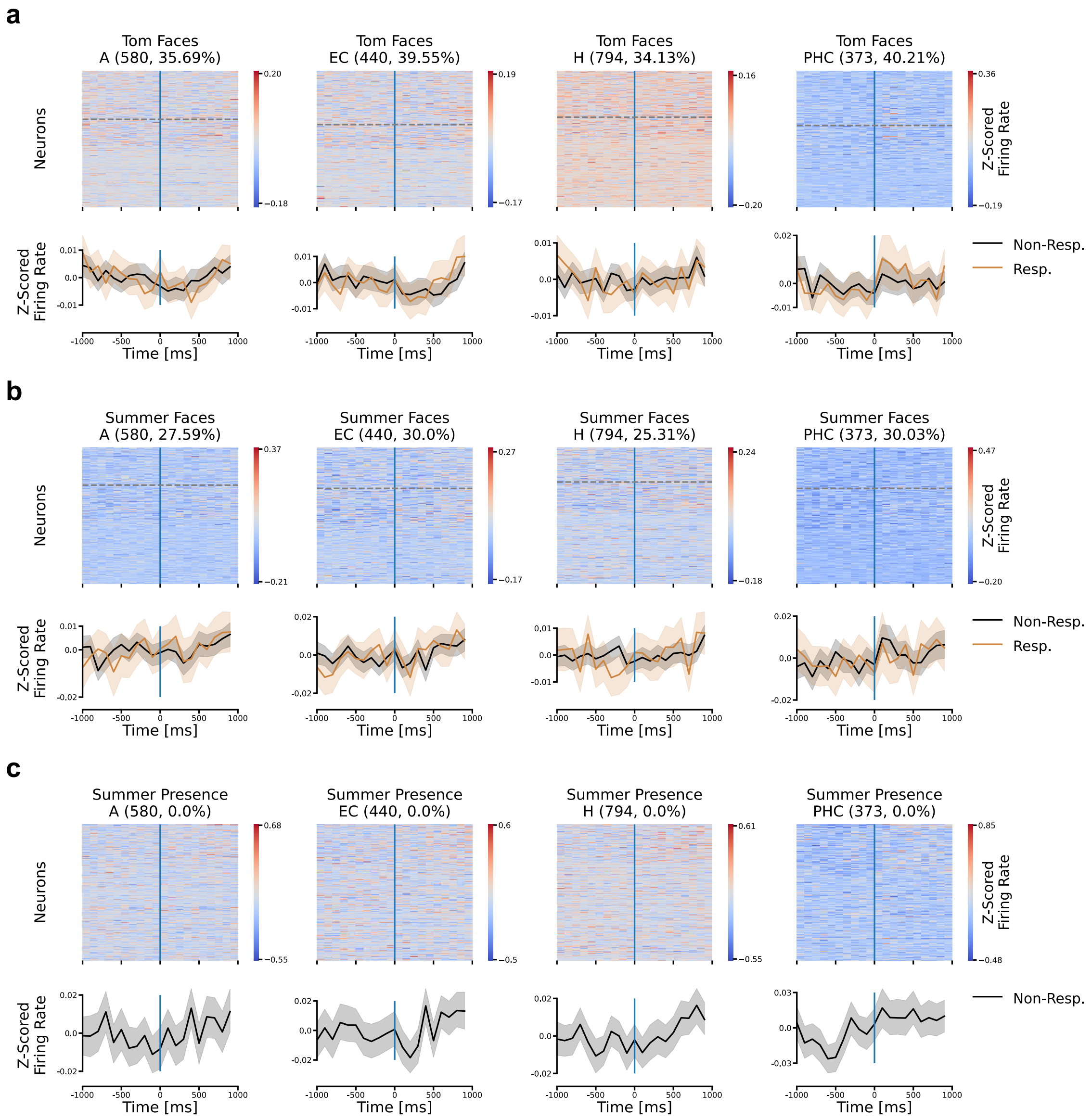

**Figure S5.** Single unit activity surrounding appearances of visual transitions and location labels during the movie presentation. **a)** Camera Cuts, **b)** Scene Cuts, **c)** Indoor, and **d)** Outdoor.

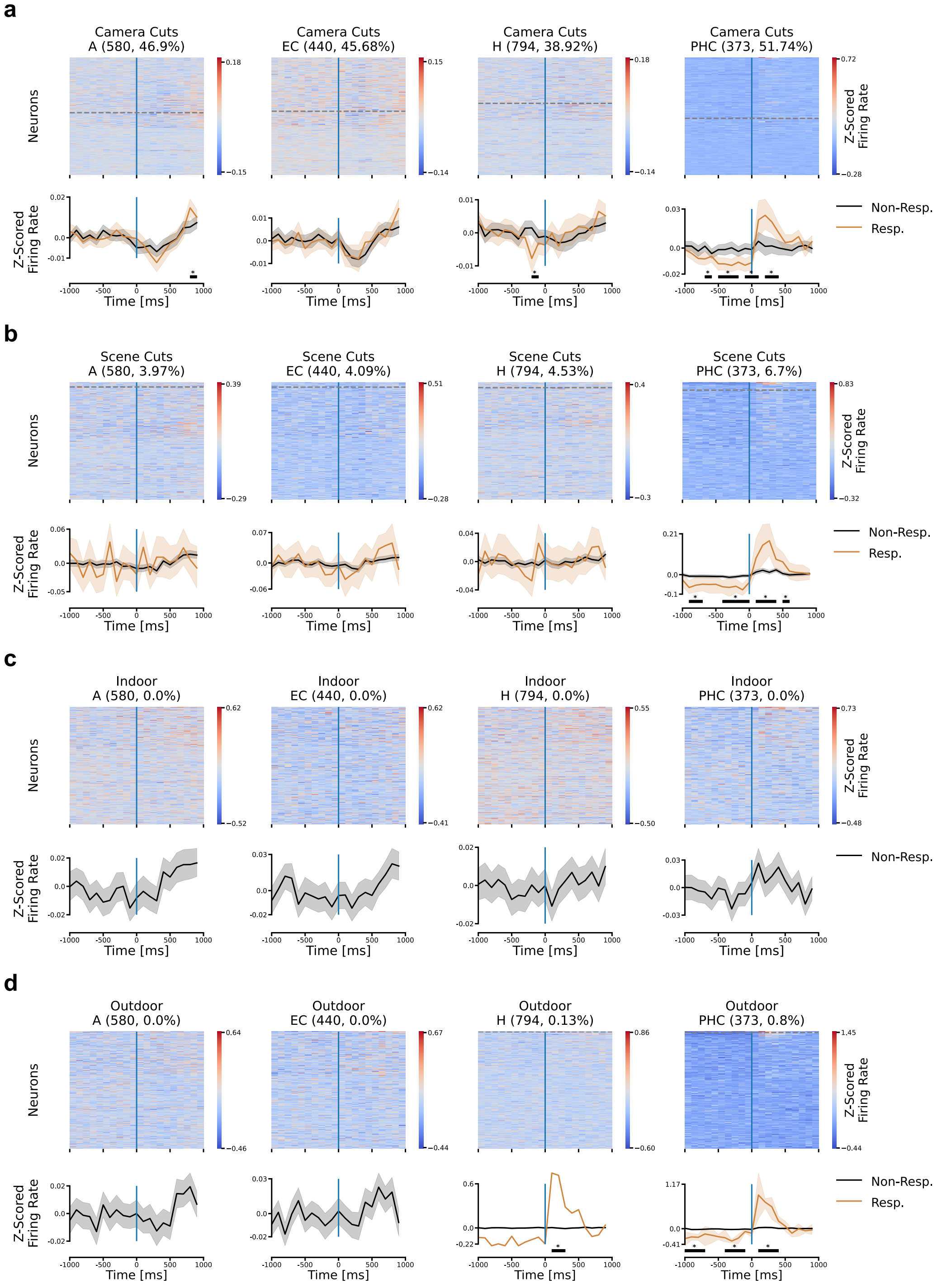

**Table S2. Overview of decoding performances for labels** Each cell contains the results of the different metrics for predicting each label from the test set. We determined the performance using 5-fold cross-validation and report the average performance across the five data splits together with the standard error of the mean. We evaluated the performance for the metrics Cohen's Kappa, F1 Score, Area Under the Precision-Recall-Curve (PR-AUC) and Area Under the Receiver Operating Characteristic Curve (AUROC). Since labels have different distributions across the movie (see Fig. 1c), chance levels vary across each labeled semantic feature and data split. Only Cohen's Kappa and AUROC have a common chance baseline across features of 0.0 resp. 0.5. As such, it is recommended to only compare performance between labels for these metrics. The best-performing semantic feature is marked in bold.

| | Cohen's Kappa | F1 Score | PR-AUC | AUROC |
|---|---|---|---|---|
| Summer | $0.3086 \pm 0.055$ | $0.651 \pm 0.02109$ | $0.6956 \pm 0.0443$ | $0.7089 \pm 0.0393$ |
| Tom | $0.23 \pm 0.0604$ | $0.3917 \pm 0.0416$ | $0.3982 \pm 0.0481$ | $0.6673 \pm 0.0366$ |
| McKenzie | $0.3162 \pm 0.0815$ | $0.3661 \pm 0.0776$ | $0.3967 \pm 0.0907$ | $0.8421 \pm 0.0359$ |
| Persons | $\mathbf{0.3599 \pm 0.0477}$ | $0.3924 \pm 0.0443$ | $0.3973 \pm 0.0551$ | $\mathbf{0.8461 \pm 0.0275}$ |
| Tom Faces | $0.2685 \pm 0.0667$ | $0.4821 \pm 0.0526$ | $0.5509 \pm 0.0578$ | $0.6654 \pm 0.0426$ |
| Summer Faces | $0.3341 \pm 0.0566$ | $0.5841 \pm 0.0431$ | $0.6116 \pm 0.0548$ | $0.7315 \pm 0.0337$ |
| Summer Presence | $0.2709 \pm 0.0445$ | $0.508 \pm 0.0346$ | $0.5418 \pm 0.0532$ | $0.6646 \pm 0.0404$ |
| Scene Cuts | $0.2007 \pm 0.0321$ | $0.2065 \pm 0.0332$ | $0.1994 \pm 0.0362$ | $0.7963 \pm 0.0619$ |
| Camera Cuts | $0.1833 \pm 0.0149$ | $0.2841 \pm 0.0201$ | $0.2374 \pm 0.0151$ | $0.6587 \pm 0.0211$ |
| Indoor/Outdoor | $0.3115 \pm 0.0597$ | $0.5106 \pm 0.039$ | $0.5709 \pm 0.0797$ | $0.7228 \pm 0.0488$ |

#### Decoding performances on the test set

We evaluated the trained decoding network on the held-out test data. We report Cohen's Kappa in the main text as this metric enables a direct comparison of decoding performance across labels despite their differing distributions within the movie. In addition, we assessed performance using the following metrics: F1 Score, Area Under the Precision-Recall-Curve (PR-AUC) and Area Under the Receiver Operating Characteristic Curve (AUROC) (see Methods for more details). Similar to Cohen's Kappa, which has a common chance baseline at 0.0, the AUROC metric has a common chance baseline at 0.5. F1 Score and PR-AUC have variable chance baselines depending on the distribution of each label. For the metrics with a common baseline (Cohen's Kappa and AUROC), performance can be directly compared, with the best performance across all labels achieved by decoding the Persons label.

#### Decoding from individual neurons

To investigate whether semantic features can be decoded from the activity of individual neurons, as opposed to populations of neurons, we chose 14 of the identified responsive neurons for Summer and evaluated their decoding performance (Fig. S6). We used a recurrence-based decoding network similar to that used for the population responses, with modifications only to the input layer. The selected neurons are spread among both patients and regions.

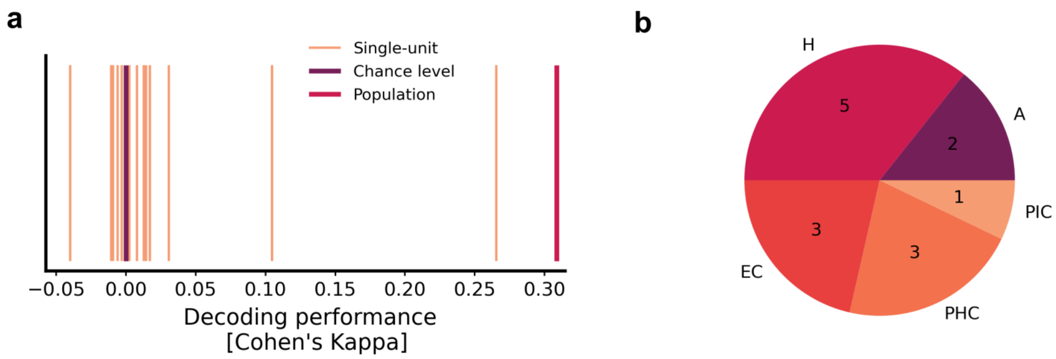


**Figure S6. Comparison of decoding performances obtained by the decoding network for population-wise and single-unit responses.** Prediction performance from 14 randomly chosen responsive single neurons for the label Summer. **a)** We compared the decoding performance for the character Summer achieved by the whole-population LSTM-based network (*Population*) to performances from models which each only took an individual neuron as input. No single-neuron model achieved significant decoding performance, even when employing a recurrence-based decoding network similar to that used for population decoding performances. **b)** The selection of single neurons is distributed across regions. The neurons collectively represent 13 different patients and are distributed across regions in the MTL according to the relative size of the whole population.

#### Data Splits: Original setup with five splits

Due to the dynamic nature of the stimuli, we used 5-fold cross-validation to assess decoding performances. The data were divided into training, validation, and test sets, with each split assigning different subsets of the data to these three categories. Importantly, all splits implement a temporal gap of at least 32 seconds between data samples belonging to different sets. The five different splits of the data are visualized in Fig. S7.

#### Data Splits: Extended setup with 20 splits

To ensure there was no overlap between training and test data in the analysis described in Fig. 7, we refined the data splitting process by implementing a nested cross-validation procedure with 20 splits instead of the original 5. Each of the five original splits was further subdivided into four splits, all of which shared the same test dataset. A representation of such a group of four splits is given in Fig. S8.

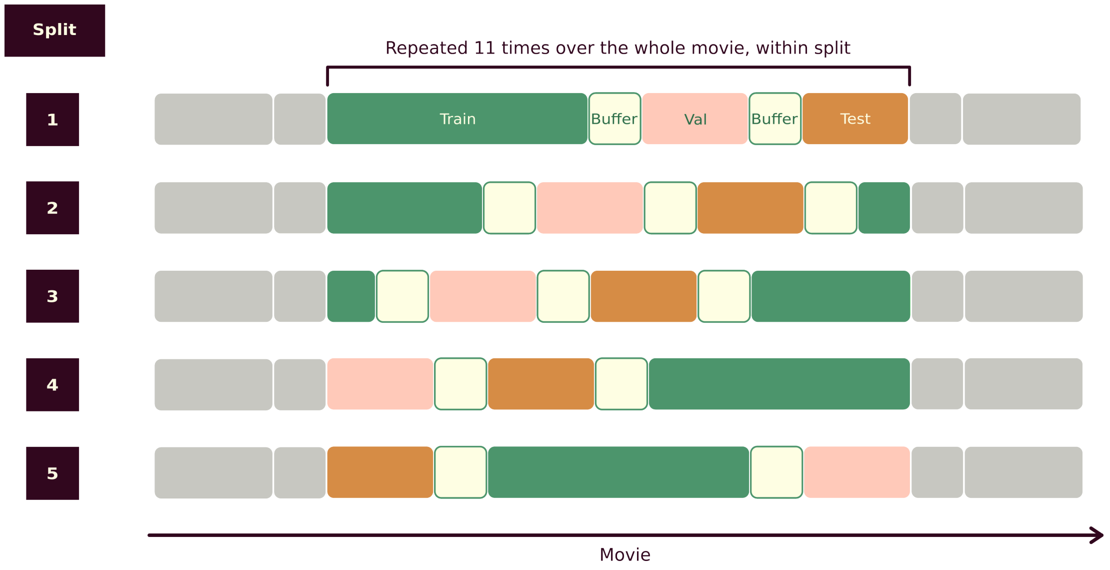


**Figure S7.** Schematic of the 5-fold cross-validation splits. The colored cells represent a common section of the movies, and each row represents a different split, or a different way of slicing the dataset into training, validation, and test sets. A buffer of 32 seconds separates each subset of samples.

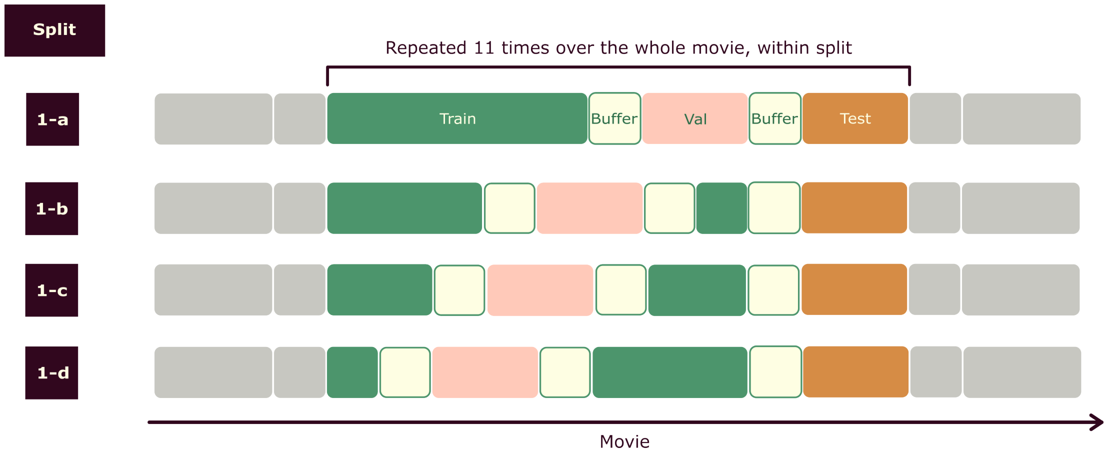


**Figure S8.** Schematic of modified splits used in conjunction with neuron rankings based on logistic regression coefficients. A set of four sub-splits sharing a common test set is shown In total, there were twenty data splits, constituting five instances of sets of four sub-splits that share a common test set, resembling the one presented here. Comparable to Fig. S7, the colored cells represent a common section of the movie, and each row represents a different division of data, i.e., a different way of slicing the dataset into training, validation, and test sets. A buffer of 32 seconds separated each subset of samples.

#### Avoiding autocorrelation in the dynamical setting

Due to the high degree of autocorrelation within the data, caused by autocorrelation within the neuronal responses and in the stimulus, it is particularly important to carefully split the data. While splitting the data, we observed that correlation within the data samples had a drastic effect on the performance. In order to obtain reliable performance on the test set, we found that two measures were critical for the implementation of the data split. First, as mentioned above, we used 5-fold cross-validation (see Methods for details). As above, for each of the five splits we subdivided the data into train, validation, and test and aimed for a distributed sampling over the whole course of the movie. The second measure introduced the concept of temporal gaps. For this, let us first assess the simplest way of splitting the data, namely randomly subdividing it intro train, validation, and test. As a consequence of random subdivision, there exist data samples in the training and test sets that lie very close in time and are therefore highly correlated. As expected, since the test data in this case is not independent of the training data, we observe remarkably high decoding performance on the test set of 0.985 (Cohen's Kappa) (Fig. S9a). In order to avoid artificially increasing the performance, we subdivided the dataset into train, validation, and test by subsampling consecutive segments of the dataset and introduced gaps between each segment. The amount of data excluded is specified by the temporal gap size. We experimented with buffer lengths ranging between 0 s (random split) and 44 s (Fig. S9a). We observed that the decoding performance converged to a constant performance of 0.26 for the label Summer for temporal gaps greater or equal to 32 s. All reported decoding results for all labels rely on data splits that have been sampled using the most conservative buffer length, 32 s. For each buffer length, we evaluated the number of overlapping scenes between training and test splits (Fig. S9b). In the movie stimuli, the average length of a scene is 36.15 s. This aligns with the presented evaluation of overlap in scenes between training and test data, which decreased towards no overlap for temporal gaps greater than 32 s. Moreover, this similarly aligns with the decreasing pattern of decoding performances. This result highlights the effect of the dynamical nature of the stimuli on the data split.

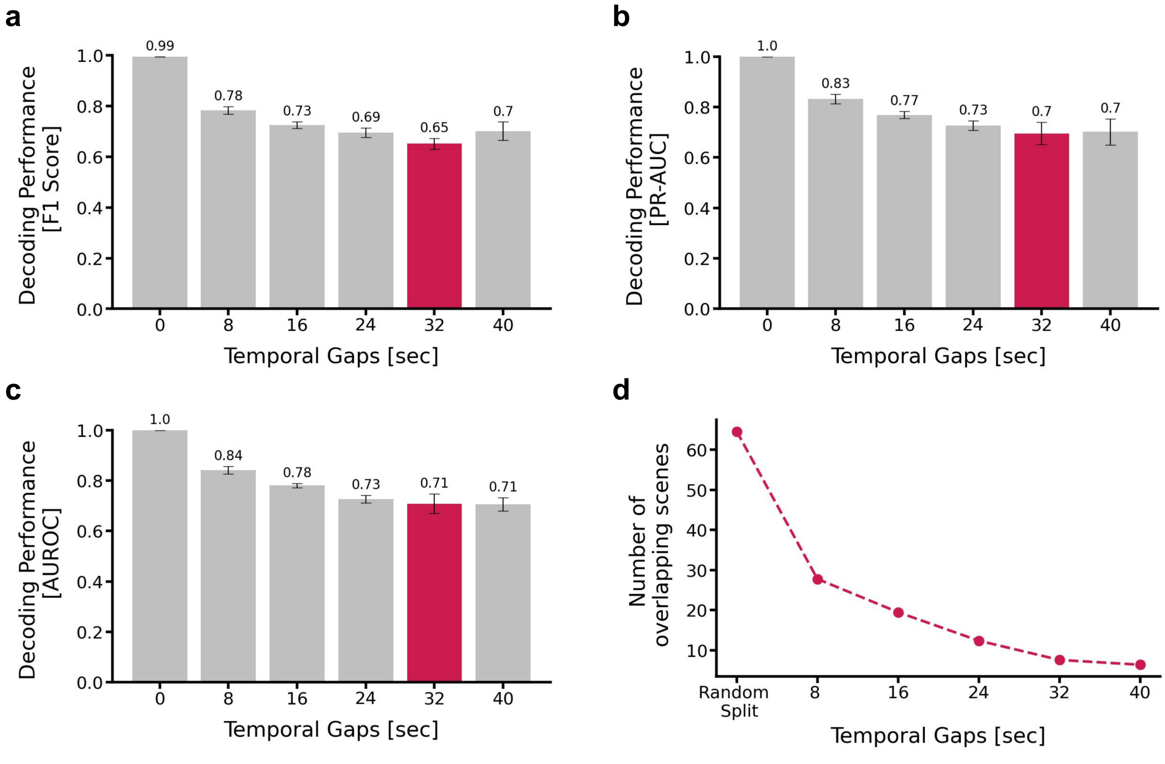


**Figure S9. Temporal Gaps between data splits reduces temporal correlation of data samples. a - c)** Effect of varying gap sizes between data splits visualized for additional metrics F1 Score, PR-AUC and AUROC evaluated on the label Summer. For all metrics, the decrease in performance stops for temporal gap sizes around 32 s. **d)** Overlap in scenes between training and test split for splits based on different gap sizes. The higher the gap size, the lower the number of overlapping scenes.

#### Decoding from responsive neurons for Tom, Scene Cuts, and Indoor/Outdoor

In Fig. 6 we evaluated decoding performances for different subpopulations based on responsive neurons or subpopulations excluding those neurons. Here, we extended the analysis to the labels Tom, Scene Cuts and Indoor/Outdoor (Fig. S10).

For the character-related label Tom, we observed that subpopulations of non-responsive neurons outperformed those containing responsive neurons for both the full population and most MTL regions. The only exception to this pattern was the entorhinal cortex. In contrast, for the visual transition label Scene Cuts, including responsive neurons resulted in improved performance compared to subpopulations without those neurons. This pattern was particularly pronounced in the parahippocampal cortex and, to a lesser extent, in the hippocampus. The Indoor/Outdoor label had only four responsive neurons in total, all from the parahippocampal cortex. Consequently, we do not report any results for the amygdala or entorhinal cortex.

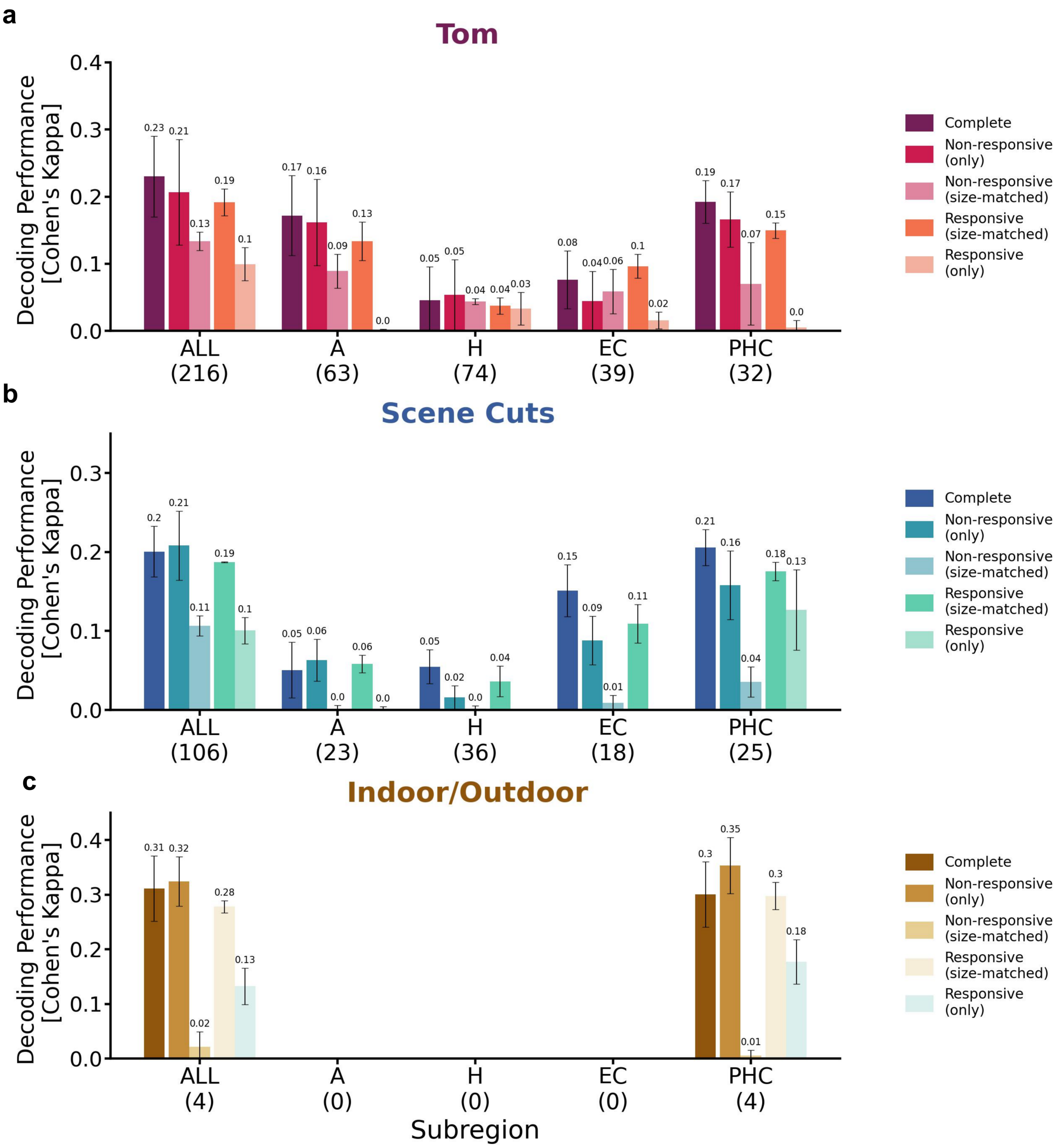


**Figure S10. Responsive neurons drive the performance for visual transitions but not characters.** Neurons identified as responsive on the single-neuron-level were utilized to restrict the population for decoding. We extended the previously discussed analysis of the character label Summer and the visual transition label Camera Cuts by the labels Tom, Scene Cuts and Indoor/Outdoor (reported performance using Cohen's Kappa, mean performance across five different data splits, error bars indicate standard error of the mean). We report the number of responsive neurons for the respective subpopulation in parentheses. **a)** Decoding performances for the label Tom evaluated across various subpopulations of neurons, divided by regions, including or excluding responsive neurons. **b) - c)** Equivalent evaluation for the labels Scene Cuts and Indoor/Outdoor.

#### Comparison of permutation results between ***Test Set Shuffle*** and ***Circular Shift***

In the main text, we evaluated the trained decoding network on a randomly shuffled version of the test data $N = 1000$ times. Chance level is given as the average of all permuted decoding performances. This approach (*Test set shuffle*) eliminates temporal dependencies within labels, leaving the input to the decoding network (the sequences of spikes) unchanged. Here, we further validate the main results of our study by applying the *Circular shift* permutation test, in which the labels are not randomly shuffled but shifted by a given amount, thereby preserving any temporal structure.

We compared the ground-truth performance against the respective null distribution for the results of Fig. 3c, for both permutation tests (Fig. S11). While the variance in the null distribution for the *Circular shift* method was more widely dispersed than for the *Test set shuffle*, the ground-truth performance consistently exceeded every permuted performance. Consistent with the *Test set shuffle* method, decoding performances for all concepts were significant at an alpha level of $0.001$ in the *Circular shift* permutation test as well.

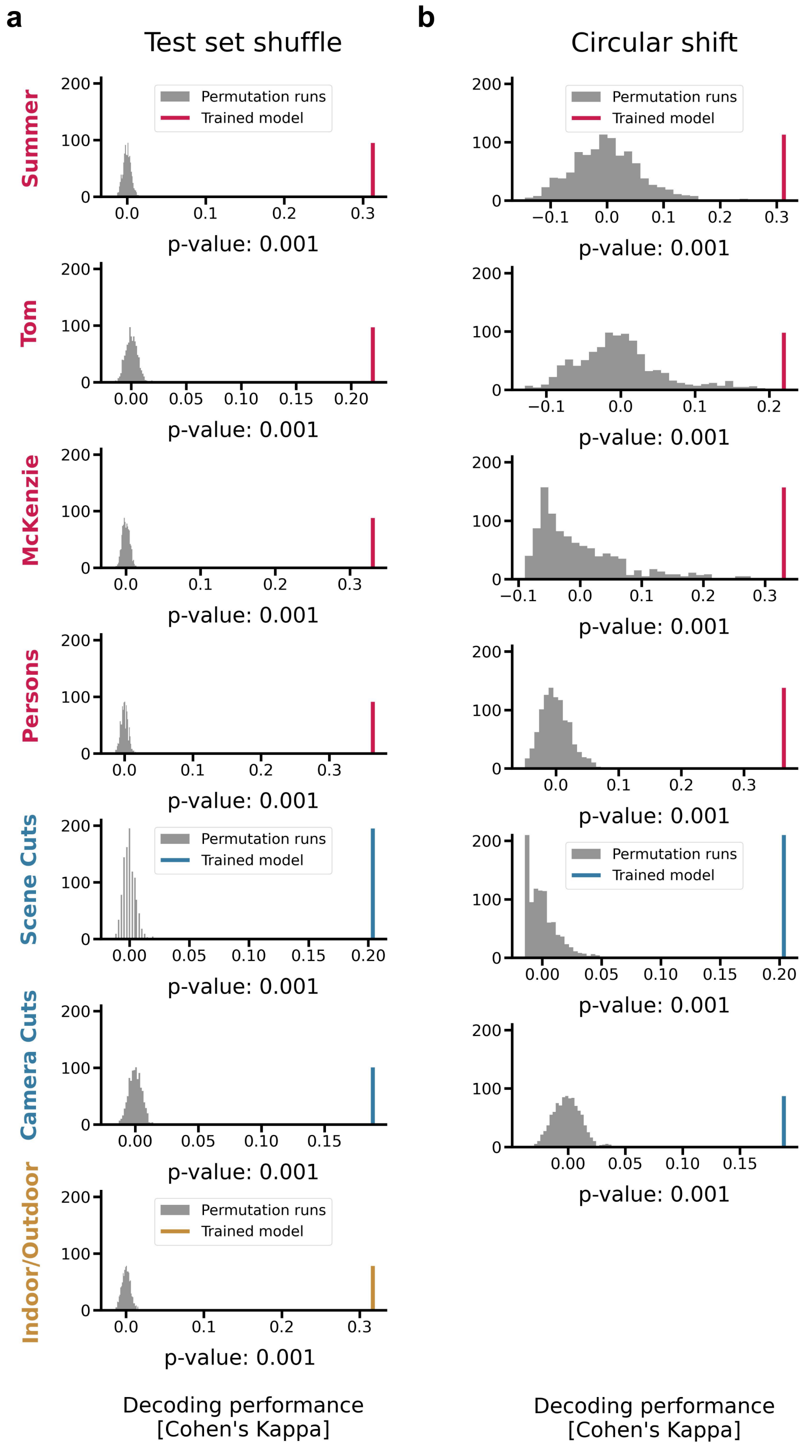


**Figure S11. Comparison of permutation results using the *Test set shuffle* and *Circular shift*.** We assessed the significance of the decoding performances through a permutation test, where trained models are assessed using permuted test labels. The resulting distribution of performances served as a null distribution which was employed to calculate a p-value for the decoding performance on ground-truth test labels (referred to as the *model's performance*). We compared the permutation results for all seven concept labels for two different ways of permutation test: **a)** A random shuffle of the test labels that destroyed the temporal structure of the concept labels (*Test set shuffle*). **b)** A circular shift of the labels that maintained the temporal structure of the concept labels (*Circular shift*). We excluded the Indoor/Outdoor label for this permutation test because it omits a certain number of frames that are neither labeled as Indoor nor Outdoor, resulting in less consistent temporal information compared to the other labels.

#### Top-ranked neurons from logistic regression decoding

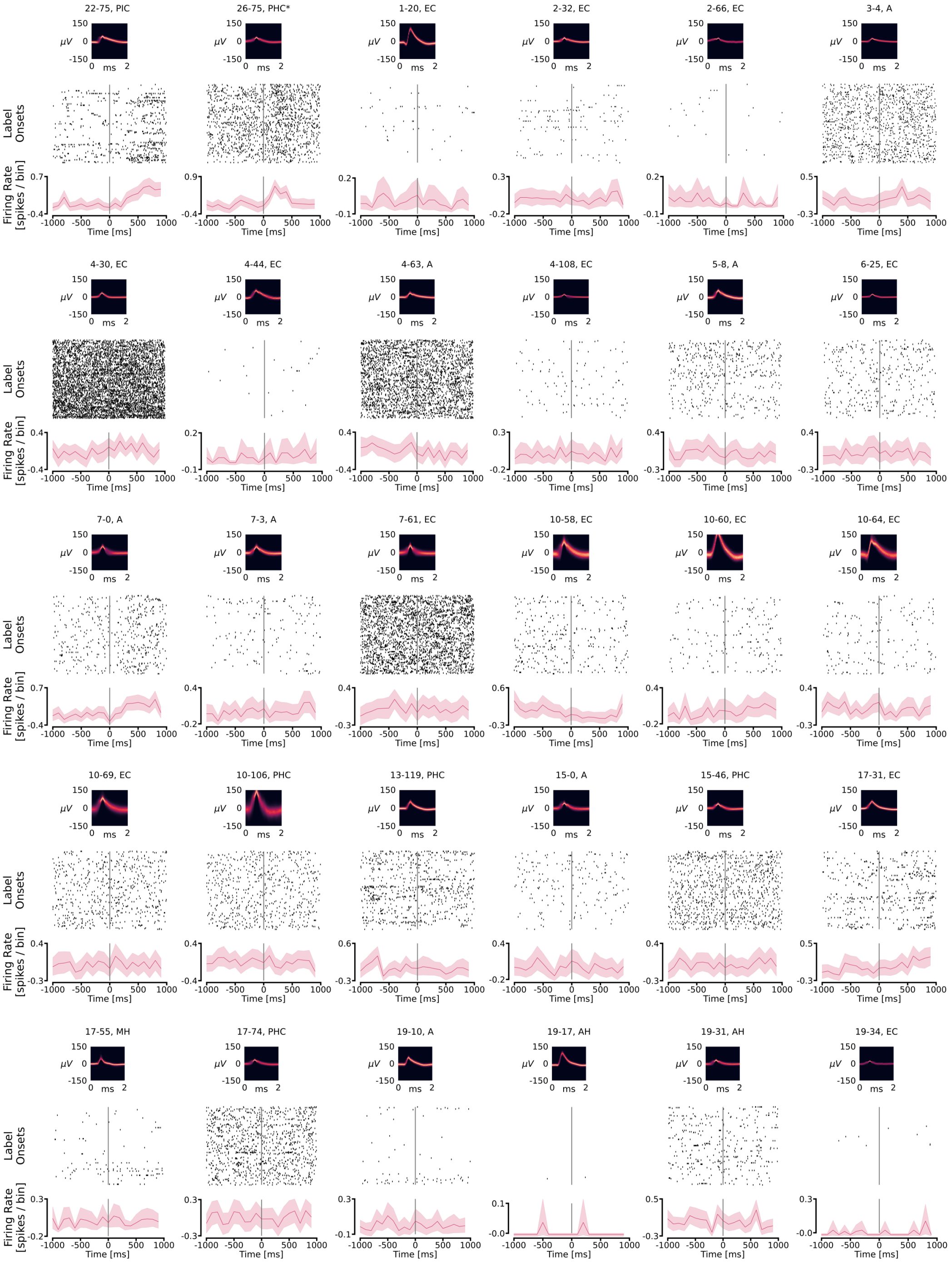


**Figure S12. Semi-random sampling of the overlapping neurons for Summer** Neurons which were separately identified as significantly responsive are denoted by *. The remaining neurons were randomly selected from the set 78 which were activated in all splits of the logistic regression training for the character label Summer.

#### Neuron ranking with non-independent training and test data

We restricted the decoding to subsets of neurons identified using the weights of a logistic regression model and implemented an extended cross-validation approach of our dataset into 20 splits. Groups of four splits shared a common test set but varied in the allocation of training and validation data (Fig. S8). From each group of four splits, we derived a ranking of neuron importance while maintaining the common test set to avoid cross-talk between training and test data.

We observed that ignoring potential cross-talk significantly distorted the decoding results, leading to artificially high performance. By restricting the analysis to 150 neurons derived from the original five splits, each with differently allocated test data, the decoding performance nearly doubled compared to decoding from the complete population of neurons, reaching 0.52%. Evaluating subsets of neurons based on rankings derived within splits sharing the same test data resulted in a significantly less pronounced effect. This underscores the need to carefully prepare the data for paradigms like ours, where both the nature of the brain activity as well as that of the dynamic, continuous stimuli induces a high degree of correlation in the dataset.

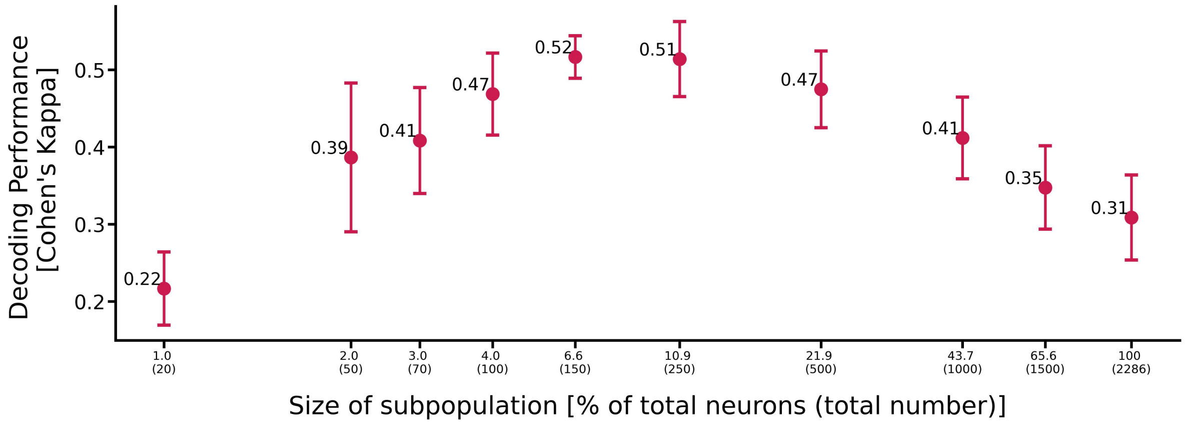


**Figure S13. Subpopulations of neurons derived with cross-talk between training and test data resulted in artificially high decoding performance.** Deriving a ranking of neurons using logistic regression weights with the original five splits can lead to data leakage between training and test data, affecting decoding performance. We evaluated decoding performances for subpopulations of top-performing neurons, where the ranking is based on the original five splits, with possible cross-talk between training and test data (sizes range from 1% to 100% of the full population, absolute numbers of neurons are reported in parenthesis). Mean performances across the splits are reported, with error bars representing the standard error of the mean. Restricting to smaller subsets of neurons leads to a significant increase in performance, with peak performance for a subpopulation of 150 neurons at 52%, nearly doubling the performance of the complete population of neurons. These results diverge significantly from those based on a clean selection of neurons, which we present by extending the dataset splitting into 20 splits, where each group of four splits shares a common set of test data.
